# Designing multiphase biomolecular condensates by coevolution of protein mixtures

**DOI:** 10.1101/2022.04.22.489187

**Authors:** Pin Yu Chew, Jerelle A. Joseph, Rosana Collepardo-Guevara, Aleks Reinhardt

**Affiliations:** Yusuf Hamied Department of Chemistry, University of Cambridge, Cambridge, CB2 1EW, United Kingdom; Department of Physics, University of Cambridge, Cambridge, CB3 0HE, United Kingdom; Department of Genetics, University of Cambridge, Cambridge, CB2 3EH, United Kingdom

**Keywords:** liquid–liquid phase separation, genetic algorithms, intrinsically disordered proteins, biomolecular condensates

## Abstract

Control of biomolecular condensates may hold considerable therapeutic potential. Intracellular condensates are highly multi-component systems in which complex phase behaviour can ensue, including the formation of architectures comprising multiple immiscible condensed phases. Conceivable avenues for manipulating condensates to bypass pathologies thus extend beyond merely controlling their stability and material properties, and relying solely on physical intuition to manipulate them is difficult because of the complexity of their composition. We address this challenge by developing an efficient computational approach to design pairs of protein sequences that result in well-separated multilayered condensates. Our method couples a genetic algorithm to a residue-resolution coarse-grained protein model. We demonstrate that we can design protein partners to form multiphase condensates containing naturally occurring proteins, such as the low-complexity domain of hnRNPA1 and its mutants, and show how homo- and heterotypic interactions must differ between proteins to result in multiphasicity.

## I. INTRODUCTION

Biomolecular condensates are involved in controlling many aspects of cell biology and pathology. By dynamically segregating particular biomolecules,^1,2^ condensates help create intracellular micro-environments that contribute to the regulation of chemical reactions^3,4^ and mediate a variety of fundamental biological processes, from cell signalling^5–9^ to RNA metabolism,^10–12^ response to stress,^13–17^ regulation of transcription^18–23^ and DNA repair.^24–26^ Moreover, dysregulation of biomolecular phase separation has been associated with a growing list of diseases from cancer to neurodegeneration.^27–32^ The link between aberrant liquid–liquid phase separation (LLPS) and disease highlights the desirability of developing new tools to manipulate the properties of condensates to bypass pathologies. Anti-cancer drugs have already been shown to concentrate preferentially into condensates,^33^ and some small molecules can modulate LLPS,^34^ making biomolecular condensates potential drug targets. A number of biomolecular condensates in vitro and in cells have been observed to display multiphase architectures: structural heterogeneity over mesoscopic length scales, where immiscible phases with different compositions coexist within the same liquid droplet. In vitro, multiphase droplets have been observed in various multi-component mixtures, including those of poly(PR) and RNA homopolymers,^35^ of polyR, polyK and polyU,^36^ and of prion-like and arginine-rich polypeptides, and RNA.^37^ Inside cells, a prime example is the nucleolus, which exhibits a multilayered architecture^38^ thought to be important for sequential processing of nascent rRNA transcripts.^10^ Similar internal structuring is also found in stress^13^ and P granules.^39–41^ The presence of multiple condensed phases in a single condensate may reflect different biological processes taking place in physically separated regions within the same compartment.^42,43^ Controlling the multiphase behaviour of condensates may thus provide additional avenues to exploit these compartments to our advantage; for example, selectively partitioning small molecules into distinct inner phases may have therapeutic potential. This leads to a fundamental question: Can we rationally and efficiently design multiphase condensates with controlled properties?

Intensive efforts to probe the driving forces for multiphase condensation suggest principles that can be considered in their design. Experiments have found that multiphase condensates form when there is competition for a shared binding partner.^36,37^ For instance, competing protein–protein and protein–RNA interactions can provide a regulatory mechanism for the organisation of multiphase condensates.^37^ Simulations of multi-component systems^44,45^ have further revealed that the phase boundary for demixed phases is sensitive to the variance of intermolecular interaction strengths: if it is sufficiently large, multiple distinct phases can form.^45^ From the physicochemical point of view, the various forms of structuring and morphological patterns of multiphase condensates have been suggested to be modulated by the difference in the interfacial free-energy densities of the phases,^38,46–49^ which in turn depend on the sequence-encoded molecular interactions of the components.^35–37,48^ Phases with high interfacial free-energy densities are expected to be engulfed by those with lower ones,^50^ while some phases form completely separate droplets due to high interfacial tensions.^51^

Although one could in principle speculate, based on physical intuition, which pairs of protein sequences might give rise to multiphase architectures, this strategy is likely only feasible for simple amino-acid sequences. Even then, the phase behaviour of multi-component systems with multiple coexisting condensed phases is far more challenging to predict than that of single-component condensates, especially given the complexity and diversity of biologically relevant proteins. Computational approaches, such as genetic algorithms, can help explore the vast size of protein sequence space by automating the design of protein sequence mutations. Broadly speaking, genetic algorithms use mechanisms inspired by biological evolution, such as crossovers and mutations, to optimise properties of a system,^52–54^ and have long been used in optimisation problems in many fields,^55–64^ including those with biological applications such as protein engineering^65–67^ and drug design.^68,69^ Genetic algorithms have recently been applied to evolve protein sequences to (de)stabilise their condensates.^70,71^ However, in order to use genetic algorithms in the context of LLPS, we first need to quantify the protein properties we wish to optimise. Recently, computer simulations have connected features of individual biomolecules to their phase behaviour.^72^ Depending on the question being addressed, models from atomistic^73–79^ via residue-level^80–86^ to minimal,^79,87–89^ alongside other computational approaches such as predictive algorithms and machine-learning methods,^84–86,90–93^ have all been used with success.

Motivated by these ideas, here we develop an evolutionary algorithm that goes beyond manipulating the stability of condensates and allows us to enforce or inhibit a desired spatial organisation of biomolecules inside multi-component condensates. We couple molecular-dynamics simulations of a residue-resolution coarse-grained protein model that achieves near quantitative agreement with experiments^83^ with a genetic algorithm^70^ to evolve protein sequences towards increasing ‘multiphasicity’, which we define to be the difference in compositions of the two coexisting phases of a multiphase condensate. The multiphasicity of a condensate increases with the purity of the two coexisting phases. We first demonstrate that we can increase the multiphasicity of a protein mixture using a genetic algorithm with an appropriate fitness function to evolve the amino-acid sequence of one of the two proteins (Sec. II A). We then show that we can design a protein sequence to act as a multiphase partner for some protein of choice by coevolution (Sec. II B), including proteins of biological relevance such as the low-complexity domain (LCD) of the heterogeneous nuclear ribonucleoprotein A1 (hnRNPA1). Finally, we analyse the changes in interaction energies (Sec. II C) and amino-acid patterning (Sec. II D) to probe the factors driving the formation of multilayered condensates.

## II. RESULTS

### A. Genetic algorithms can improve separation of two-protein multilayered systems

Genetic algorithms can effectively evolve amino-acid sequences to find mutations that give rise to desired changes in phase behaviour.^70,71^ During the evolution, each protein in a population is associated with a fitness value, as computed with a fitness function, and the population is evolved with local mutations and crossovers between individual sequences towards a fitter – albeit not necessarily the fittest – solution. Rather than determining an optimal solution (however defined), the genetic algorithm follows local gradients in fitness space towards a better solution that is not drastically dissimilar to the starting sequence. The success and efficiency of the genetic algorithm depend on the quality of the fitness function, which should be relatively simple and inexpensive to compute, while at the same time sufficiently complex to act as a suitable proxy for the property being evolved. For simplicity, here we focus on evolving two-component protein condensates towards higher multiphasicity (as defined above). Nonetheless, our approach can readily be generalised to condensates with a larger number of components, which are in any case expected to behave similarly.^45^

To design our fitness function, we start with the simplest metric of multiphasicity we could conceive for a two-component system: the difference in the number densities between the two different proteins at the centre of the multilayered condensate. This quantity is small when the two-component mixture phase-separates into a homogeneous condensate (i.e. low multiphasicity) and large when it phase-separates into a multilayered condensate with each layer enriched in a different protein (i.e. high multiphasicity) [Fig. 1**a**]. However, if the two-component mixture phase-separates into a homogeneous condensate that is depleted of one of the two proteins and a dilute phase that is enriched in the depleted protein, this metric is large even though the multiphasicity is low [Fig. 1**a(iii)**]. To avoid this, we introduce a second term in the fitness function that penalises the accumulation of either protein in the dilute phase, namely the sum of the number densities of the two components far away from the condensate [Eq. (1)], scaled by a weighting parameter *s*. A large *s* might seem desirable to ensure the enrichment of both components inside the condensate; however, when its value is too large, it dominates the fitness function, rendering the first term irrelevant in magnitude, and can actually favour homogeneous condensates instead. In general, this weighting parameter can be tuned as necessary depending on the specific system of interest [see Fig. S6].

**Figure 1.**
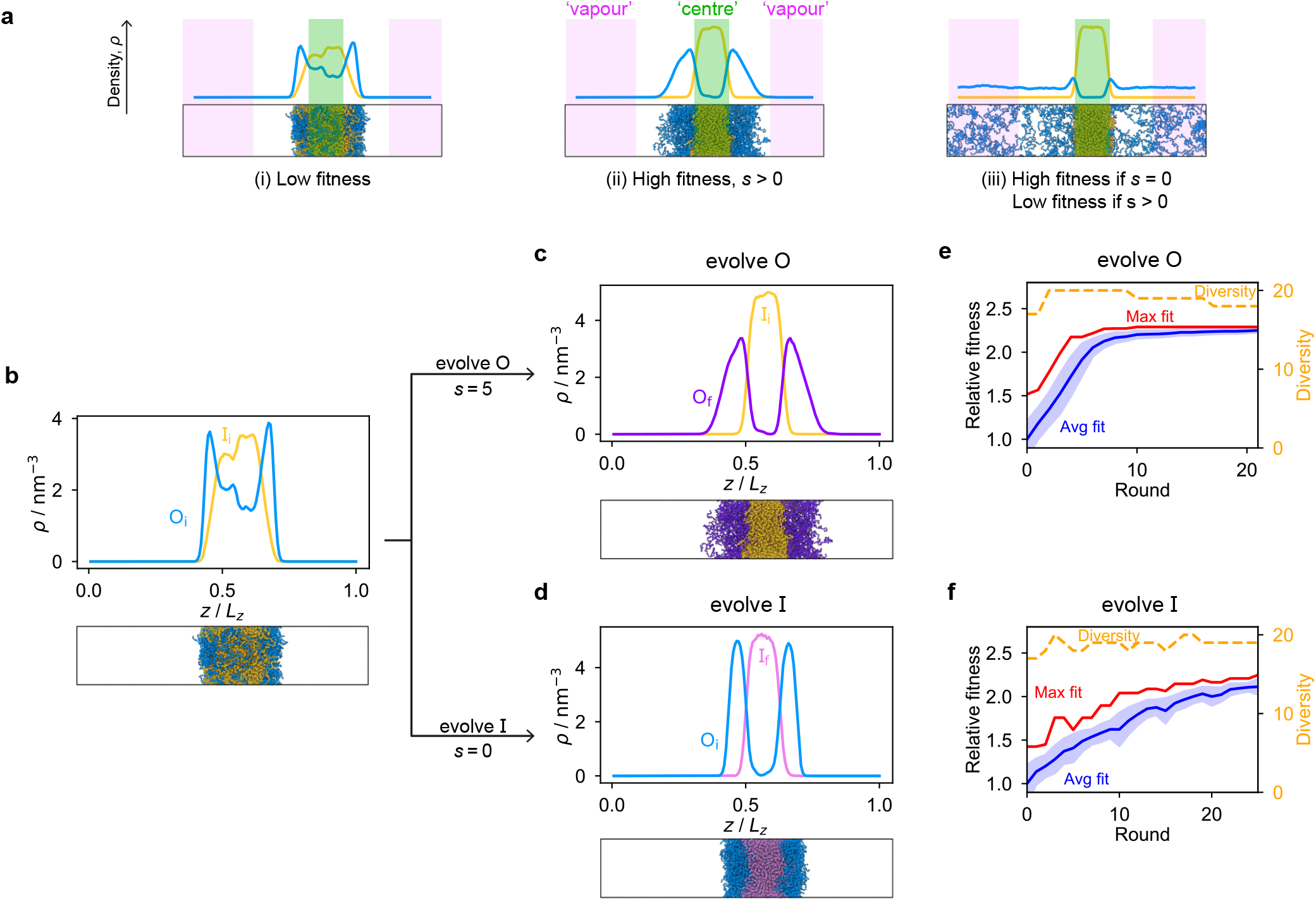
The genetic algorithm can improve multiphasicity. **a** Examples of systems with high and low fitness values. The parameter *s* determines the trade-off between increasing the difference in compositions between the two phases and obtaining two stable liquid-like phases alongside a vapour-like phase. **b** Density profile (number density *ρ* against the long axis of the simulation box) of the initial two-component system considered. Protein I is enriched in the inner layer and protein O in the outer layer. Here O_i_ = (FAFAA)_10_ and I_i_ = F_50_ with random noise added by introducing mutations with probability 0.60 to the latter such that both sequences mix to give a system with low multiphasicity. **c** Density profile of the final evolved system with maximum fitness when protein O is evolved, and **d** when protein I is evolved in separate runs. Snapshots of the corresponding multiphase droplets are provided for each case. **e, f** Fitness (relative to the average fitness in round 0) as a function of genetic-algorithm progression. The shaded area corresponds to the standard deviation of the fitness across the population at each round. In both cases, a high population diversity is maintained.

When evolving a two-component protein system towards increasing multiphasicity, the goal is to obtain a set of mutations to the amino-acid sequences of the two proteins such that the mutated proteins form a condensate with a more segregated multilayered architecture than the starting pair. We refer to the protein enriched in the core of the multilayered condensate as the ‘inner protein’ or ‘protein I’, and the one concentrated in the outer layer as the ‘outer protein’ or ‘protein O’. There are several routes one could take: one could evolve either the inner or the outer sequence in separate evolution runs, or even evolve both sequences, simultaneously or alternately, in the same run. In our evolution runs, we evolve either the inner sequence or the outer sequence whilst keeping the other sequence unchanged in order to simplify the subsequent analysis of driving forces. To evolve the sequence, we apply the genetic algorithm as described above (and in the Methods section), computing a sequence’s fitness by performing direct-coexistence moleculardynamics simulations of the two-component mixture using our residue-resolution coarse-grained model, Mpipi,^83^ at a fixed temperature.

As a preliminary test of our approach, we first consider a mixture of (FAFAA)_10_ and F_50_ and add sufficient mutations (i.e. random noise) to the latter to ensure the initial condensate has a low fitness. The mutations are added in a similar way to how the initial population of sequences is generated in the genetic algorithm (Methods), but with a replacement probability of 0.60. In this initial state, (FAFAA)_10_ is slightly enriched at the interface and is deemed the outer sequence, but there is an appreciable degree of mixing with the inner sequence. We then perform two separate evolution runs, evolving (a) the inner sequence whilst keeping the outer unchanged, and (b) the outer sequence whilst keeping the inner sequence unchanged. A comparison of the density profiles of the initial system to that of the final systems with the maximum fitness [Fig. 1**b–d**] confirms that our fitness function can successfully guide the initial system towards increasing multiphasicity in both cases. When the inner sequence is evolved, a stable multilayered condensate can be obtained with a weighting parameter *s* = 0 used in the fitness function; by contrast, when evolving the outer sequence, the final result is sensitive to the value of the parameter *s* and *s* > 0 must be used. We show in Fig. S6 the final evolved systems obtained with different values of *s*. Additionally, the genetic-algorithm progressions in each case are depicted [Fig. 1**e,f**]. We plot the mean fitness of the population (blue curve), the fitness of the fittest individual (red curve) and the number of distinct sequences in the population (i.e. its ‘diversity’; yellow curve).

### B. Multilayered condensates can be designed by coevolving a partner protein sequence alongside a protein of interest

Armed with an effective fitness function for multiphasicity, we next set out to use our genetic algorithm to guide the design of partner proteins that result in multilayered condensates alongside known phase-separating proteins of interest (e.g. a naturally occurring protein, such as hnRNPA1 LCD used below). There are many challenges involved in designing a partner protein for this purpose, such as ensuring that it phase-separates at similar experimental conditions as the protein of interest (e.g. salt, pH, temperature), and that it establishes suitable associative interactions with the protein of interest to form a single multilayered condensate. Thus, to facilitate convergence in this more complex scenario, we start our coevolution approach [Fig. 2**a**] from an initial reference system of two proteins, both different from our protein of interest, which phase separates into a multilayered structure with a high degree of multiphasicity. The initial reference systems used in the coevolution runs were designed using simple generic sequences of amino acids based on knowledge from previous experimental and theoretical studies showing that immiscible phases form when interaction strengths between their components are sufficiently different.^35,36,45,48,49,94^ To arrive at our protein of interest, we systematically mutate one of the two reference sequences throughout the coevolution run. These systematic mutations are done gradually, once every 5 rounds, by randomly changing ∼10 % of the residues of this protein selected from those that have yet to be changed. Simultaneously, we evolve the other sequence with the genetic algorithm using our fitness function [Eq. (1)].

**Figure 2.**
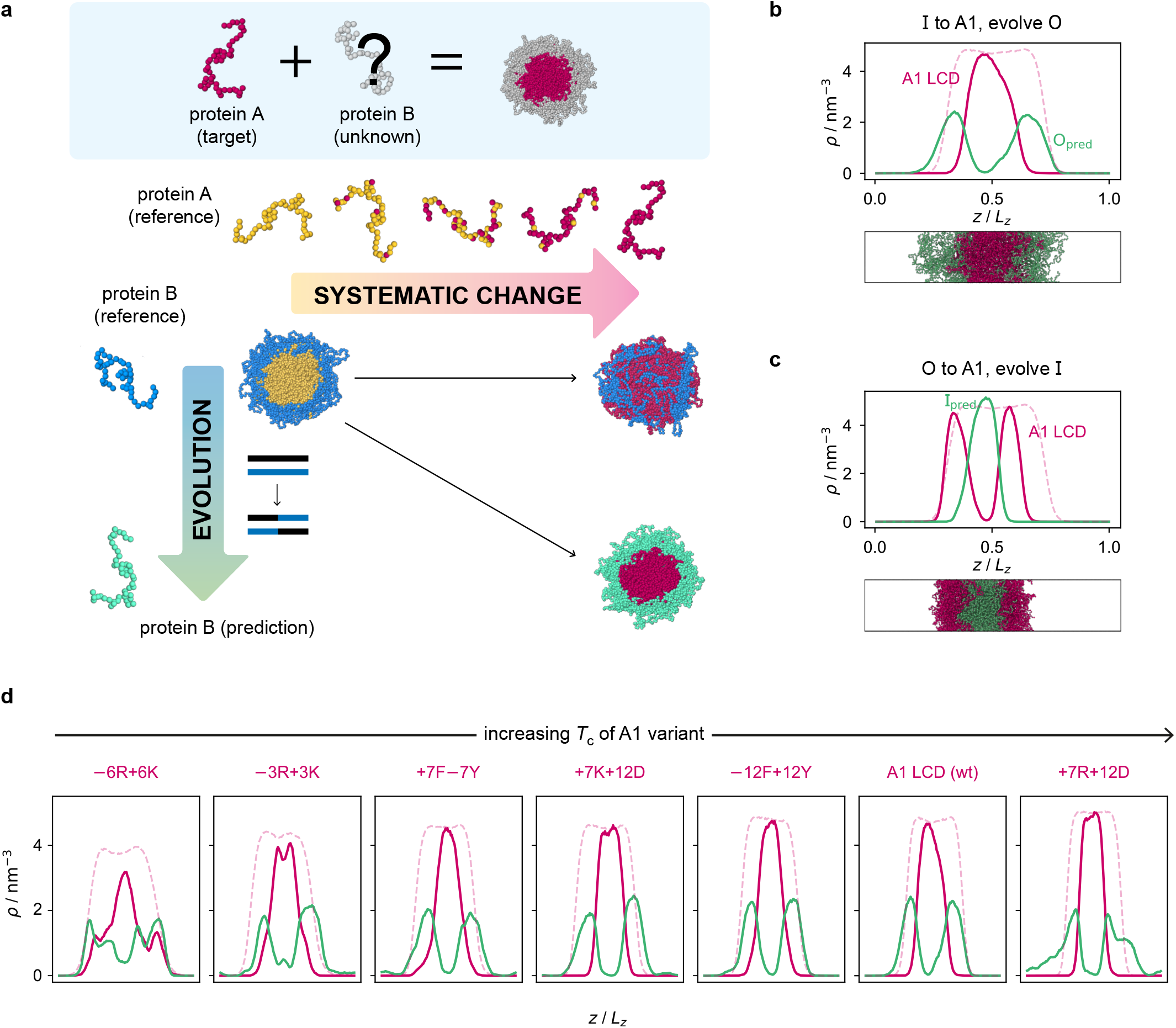
Coevolution of a multilayered condensate partner for A1 LCD. **a** Illustration of the coevolution approach. Starting from an initial two-component reference system with high multiphasicity, we systematically change one sequence to the predetermined target protein (yellow to pink) while simultaneously evolving the other sequence (blue to green) to predict a partner sequence that forms a multilayered condensate with the target protein. **b** Density profile of the final evolved system with maximum fitness from the coevolution run where A1 LCD is designed to be at the centre and **c** where A1 LCD is designed to be on the outside of the condensate. The pink dashed line is the density profile of a single-component system of A1 LCD equilibrated at the same temperature (250 K). **d** Density profiles of the final evolved systems with maximum fitness from the coevolution runs with different variants of A1 LCD as the inner sequence. The densities of the A1 variant and the predicted outer partner sequence are plotted in pink and green, respectively. The pink dashed lines are the density profiles of a single-component system of the corresponding A1 variant equilibrated at the same temperature. The A1 variants are arranged in order of increasing upper critical solution temperature.

Since only a small proportion of the residues in the sequence that is systematically changed are modified each time, multiphasicity can be maintained at least to some degree throughout the process. This procedure ensures that there is a gradient of the fitness in sequence space in the direction of increasing multiphasicity which the genetic algorithm can evolve towards at every round during the coevolution. If one started from the reference multilayered system and changed one of the sequences to the target sequence in one go, this may result in full mixing of the two components within a homogeneous liquid-like phase and hence the loss of multiphasicity altogether. From combinatorics, there are numerous sequences that can form a homogeneous condensate with the target sequence; using the genetic algorithm starting from a fully mixed state would therefore be inefficient, as the initial random search for possible mutations that result in multiphasicity would be slow before there is a gradient in sequence space towards increasing multiphasicity that can be exploited by the genetic algorithm.

Of course in some cases, changing one sequence in the reference system to the target protein may still give a multilayered system, albeit likely with a lower degree of multiphasicity. Alternatively, one may also be able to propose, based on physical intuition and understanding of the intermolecular interactions that give rise to the formation of multiphase condensates, a protein sequence that can form a multilayered condensate with the target protein. In such situations, it is possible to use the genetic algorithm directly to evolve the system towards an increasing degree of multiphasicity, as discussed in Sec. II A. However, importantly, the coevolution approach we have outlined would find a possible solution much more efficiently even if an initial multilayered system with the target protein is unknown and difficult to predict by hand.

Here, to demonstrate the robustness of the coevolution approach, we select the initial reference systems and target sequences in the coevolution runs such that making the systematic change directly in one go results in complete mixing to give a single homogeneous liquid-like phase [cf. Fig. S19**a**]. We show the results of the coevolution approach tested on systems with simple generic sequences in Sec. S10.11.

#### A multiphasic partner can be found for hnRNPA1 LCD and its variants

To demonstrate how the coevolution approach can be used to design multiphase condensates containing naturally occurring phase-separating proteins, we turn our attention to the LCD of hnRNPA1 (denoted here as A1 LCD). We first focus on designing a multilayered condensate with A1 LCD concentrated at the centre, and below we look at the converse case. To predict the partner sequence that forms a multilayered condensate with A1 LCD at the centre, we start our procedure with a mixture of I = F_135_ and O = (FAFAA)_10_, and then systematically change the inner protein to A1 LCD whilst evolving the outer protein using the genetic algorithm. We choose the initial inner protein such that its length is the same as that of A1 LCD. During the first 45 rounds of the coevolution procedure, the residues of the inner protein F_135_ are systematically and gradually changed to those of A1 LCD [Fig. S7**e**], whilst the outer protein (FAFAA)_10_ is evolved. We continue the genetic algorithm on the outer protein for an additional 20 rounds to increase the degree of multiphasicity of the system further. We show the density profiles and snapshots of the initial system and the final evolved system at the end of the coevolution run in Fig. S7**b,c**; the final system exhibits two liquid-like phases of different composition with A1 LCD enriched in the centre, demonstrating that the coevolution approach is able to predict a partner sequence that forms a multilayered condensate with A1 LCD.

The genetic-algorithm progression of this coevolution run is shown in Fig. S7**d**. The average fitness and the maximum fitness suddenly decrease at rounds where systematic changes are made and the fitness of the entire population is recalculated. Although the outer sequence is evolved using the genetic algorithm for a considerable number of rounds after the inner sequence has been completely changed to A1 LCD, the maximum fitness does not improve in these rounds, suggesting that we have reached a local maximum in the fitness function. Changing the starting sequence of the protein being evolved does not appear to offer sufficient flexibility to support a higher degree of multiphasicity. We hypothesise that a higher multiphasicity might be achieved by increasing the length of the partner sequence that is evolved, since more sequence variations are possible with a longer sequence. To test this idea, we increase the length of the outer protein from 50 to 100 residues, but keeping the total number of protein residues unchanged, and repeat our coevolution procedure. Specifically, we start the coevolution from a mixture of I = F_135_ and O = (FAFAA)_20_, and then systematically and gradually change the inner protein to A1 LCD whilst evolving the outer sequence [Fig. S9**a**]. The density profile of the final evolved system with the longer partner sequence is shown in Fig. 2**b** [see also Fig. S11**b** and Fig. S10**a**]. As hypothesised, the degree of multiphasicity of the final system is considerably improved with a longer protein partner, likely, as speculated, because of the greater flexibility in sequence choice with a longer protein.

In principle, the resulting partner sequences obtained from the coevolution run depend on the identity of the two proteins in the initial reference system, and it is not immediately obvious how to choose the reference system sensibly. Indeed, our work highlights that the solution to this problem is not unique and multiple different partner sequences can form diverse multilayered condensates with a specific target protein of interest. If we wish to find a possible solution, rather than a specific one, starting from any reference multilayered system should be feasible. To demonstrate this, we repeat the coevolution run starting from a different reference system with different protein sequences, namely a mixture of O = N_100_ and I = Y_135_. The latter, at the centre of the multilayered condensate, is then systematically changed to A1 LCD, while the outer protein is evolved using the genetic algorithm. Density profiles [Fig. S8**b,c**] confirm that the coevolution approach is again successful; of course, unsurprisingly, the final evolved partner sequence is considerably different from before, since we expect it to retain at least some features from the initial reference sequence.

We now test the ability of our coevolution algorithm to predict the amino-acid sequence of a protein partner that forms a multilayered condensates with A1 LCD concentrated towards the interface of the condensate. To do so, we construct a reference multilayered system of I = (FAFAA)_20_ and O = (FIQII)_27_, but now we change the outer protein systematically and gradually to A1 LCD whilst evolving the inner one. As desired, we show [Fig. 2**c**] that the system forms a multilayered condensate with A1 LCD towards the interface [see also Fig. S11**c** and Fig. S10**b** for further details].

Finally, to demonstrate the robustness of the approach to the target protein sequence, and to allow us to investigate if there are any overarching governing principles of multiphasicity that we can identify, we repeat the coevolution approach to find partner sequences for different variants of A1 LCD. In these cases, we choose the final multilayered condensates to have the A1 LCD variant concentrated in the centre. The phase diagrams of a set of A1 LCD variants have recently been determined both experimentally^95^ and computationally using the same coarsegrained model that we have used in this work.^83^ We consider sequences with similar, higher and lower upper critical solution temperatures compared to the wild type (WT), ensuring that sequences with distinct features are represented. In particular, we focus on two aromatic variants, +7F−7Y and −12F+12Y, two mixed-charged variants, +7R+12D and +7K+12D, and two arginine–lysine variants, −6R+6K and −3R+3K. As for the WT, we start coevolution runs from a mixture of I = F_135_ and O = (FAFAA)_20_. We show the density profiles of the final evolved systems in Fig. 2**d** and the corresponding genetic-algorithm progressions in Fig. S10**c–h**. The coevolution approach successfully predicts a suitable partner sequence for each A1 variant. The overall trends in the change in fitness are similar for all of the variants, although the final systems have varying degrees of multiphasicity. In particular, the final evolved systems with the −6R+6K and −3R+3K variants, which have considerably lower critical temperatures than the WT, are less well separated. We speculate that this result can be improved with a longer partner sequence, as we have shown for the WT.

### C. Multilayered condensates are driven by difference in component interaction strengths

Predicting the partner sequence to design multilayered condensates containing a certain target sequence is of practical importance. However, molecular simulations allow us to go one step further and understand the underlying physical and molecular driving forces for the observed behaviours. To identify which properties are important for multiphasicity, we analyse the changes in the composition and patterning of the evolved sequence in the evolution runs, where one component remained unchanged throughout, and in the coevolution runs with systematic changes of one component to A1 LCD.

We summarise the changes in amino-acid composition of the evolved sequences in Fig. 3. In the evolution runs where the outer sequence is evolved while the inner sequence is kept unchanged, the amino-acid composition of the evolved sequence changes to favour fewer aromatic residues [Fig. 3**a**]. By contrast, when the inner sequence is evolved while the outer sequence is kept unchanged, evolution favours a higher number of aromatics [Fig. 3**b**]. These observations suggest that within multilayered condensates, when other features are kept constant (e.g. protein charge, disorder and length), protein sequences with a higher aromatic content are likely to cluster towards the centre. The mean interaction strengths amongst amino acids change across the evolution runs; we use the parameter *ε*_*i*,Mpipi_ of the Mpipi model [see Methods] to estimate these changes. The average of *ε*_*i*,Mpipi_ across all residues in the evolved sequence decreases over the course of the genetic-algorithm run when the outer sequence is evolved [Fig. S12**a**], but increases when the inner sequence is evolved [Fig. S12**b**]. By comparing the final evolved sequences with maximum fitness across three independent evolution runs [Fig. 3], we note that residues become more strongly interacting on average when the inner sequence is evolved, and conversely, more weakly interacting when the outer sequence is evolved. As evidenced by Fig. 3, there are numerous ways of achieving such a change in interaction strengths, and the solution to this optimisation problem is unsurprisingly not unique. This is because for a given predetermined sequence, from combinatorics, there should exist multiple different protein sequences that can form a multilayered condensate with it, i.e. many sequences are similarly fit and the well of fitness in sequence space is broad. This important observation puts forward the idea that the formation of multiphase condensates is a general phenomenon requiring only a generic set of interaction rules governed by the overall chemistry of the functional groups involved (e.g. π-rich, charged, hydrophilic), rather than highly sequence-specific features.

**Figure 3.**
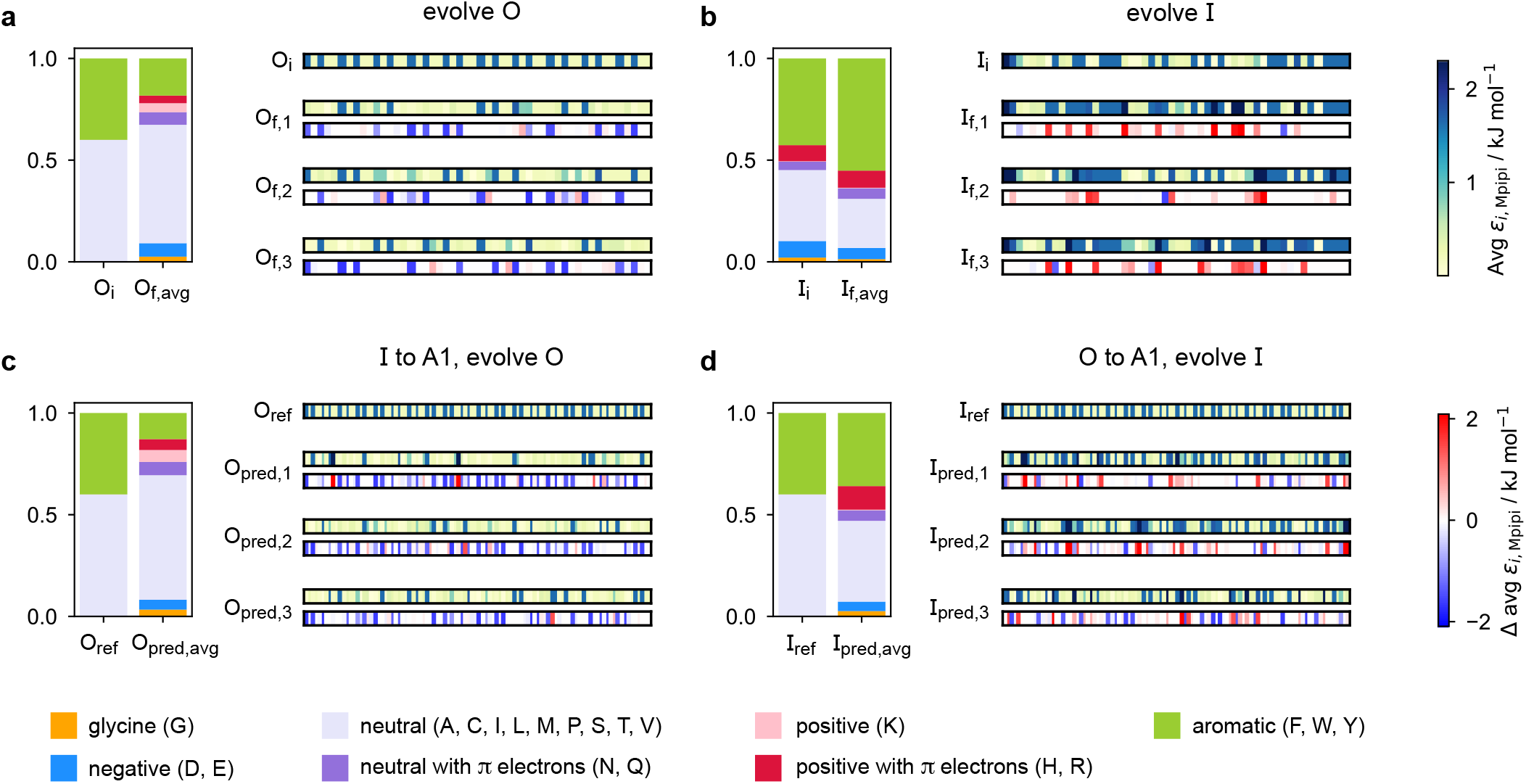
Amino-acid composition and patterning on evolution. Comparison of the amino-acid composition and sequence patterning between the initial and final evolved sequences with maximum fitness in the evolution runs where **a** the outer sequence is evolved and the inner sequence is kept unchanged and **b** the inner sequence is evolved and the outer sequence is kept unchanged; and the coevolution runs where A1 LCD is designed to be **c** the inner sequence and **d** the outer sequence in the final multilayered system. For each case, we show the composition of the initial sequence and the final evolved sequence averaged across three independent runs. To illustrate the final evolved sequences in three independent runs in each case, we plot for each residue *i* along the sequence the absolute value of and the change in *ε*_*i*,Mpipi_ compared to the residue in that position in the initial sequence. The value of *ε*_*i*,Mpipi_ estimates the interaction strength of the residue in the coarse-grained model.

When designing multiphase condensates that have the phase of A1 LCD proteins or its variants at the centre, we observe that the proportion of aromatic residues in the evolved partner protein decreases [Fig. 3**c**, Fig. S12**c**]. Consistently, the more strongly interacting residues are preferentially replaced by less strongly interacting ones throughout the evolved partner protein sequence. This change in composition of the evolved sequence is similar to the trend we observe in the evolution run where the outer sequence is evolved and the inner sequence is kept constant. The final evolved sequences in the coevolution runs with the different A1 variants are also similar in terms of the change in composition [Fig. S15**a**], even though we selected variants with different features. The main driving force for the evolution towards increasing multiphasicity of condensates with A1 LCD at the centre is the decrease in the average interaction strength of the outer sequence. However, for the case where A1 LCD is designed to be on the outside of the multilayered condensate and the inner sequence is evolved, the proportion of aromatic residues in the final evolved sequences is similar to the initial inner sequence and it is less clear whether the residues become more or less strongly interacting throughout the sequence [Fig. 3**d**, Fig. S12**d**]. This is not entirely surprising, since in the coevolution runs the two sequences in the system are being changed and evolved simultaneously, so we cannot necessarily expect the same trends as when only one sequence is evolved.

To rationalise why the compositional changes we observe in the evolved sequences favour multiphasicity, we compute the strengths of homotypic and heterotypic interactions between proteins forming the inner and the outer phases, and their changes throughout the evolution runs [Fig. 4]. Overall, our analyses reveal that formation of two-component multilayered condensates depends on three crucial requirements. First, larger differences in the strengths of homotypic interactions of the different species (i.e. inner–inner versus outer–outer) favour demixing of the components into separate phases [Fig. 4**d**]. Second, the proteins that establish the stronger homotypic interactions form the inner phase of the multilayered condensate. In other words, the inner–inner interaction is always the strongest, likely because such an arrangement guarantees saturation of binding sites that can form the most energetically favourable interactions. Third, the strength of heterotypic interactions should lie on a critical ‘sweet spot’: small enough to favour demixing into separate phases, but sufficiently large (i.e. comparable with the weaker outer–outer homotypic interactions) to keep the separate phases coexisting inside the same condensate.

**Figure 4.**
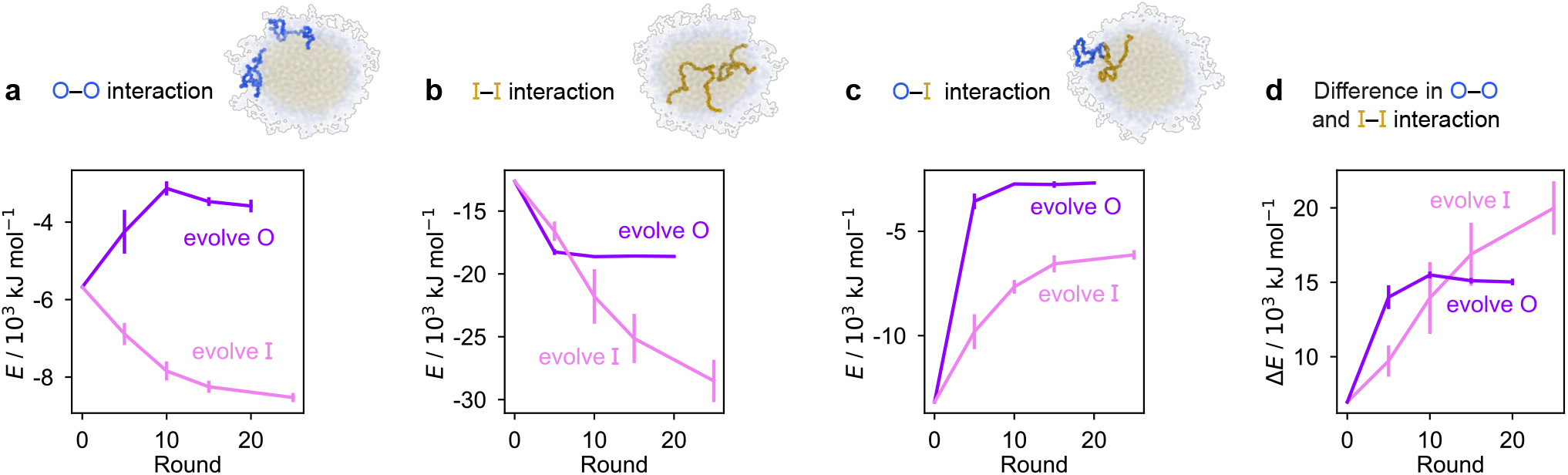
Homo- and heterotypic interaction strengths control multiphasicity. The change in the interaction energies of intermolecular homotypic interactions between proteins enriched in the **a** outer and **b** inner layers, **c** heterotypic interactions between the two proteins and **d** the difference between the outer–outer and inner–inner homotypic interactions for the system with the fittest individual. Purple curves correspond to the system where the outer protein was evolved and the inner was unchanged, and pink curves correspond to the converse. Error bars correspond to the standard deviation across three independent runs.

Our evolution and co-evolution algorithms induce changes in the homotypic and heterotypic interactions across the evolution that depend on the properties of the starting condensates and which of the proteins is being evolved. For the evolution runs where one sequence is evolved and the other kept unchanged, our starting system exhibits relatively modest strengths for both the inner–inner and outer–outer homotypic interactions, and comparatively strong heterotypic interactions. These heterotypic interactions become weaker throughout the evolution runs when either the inner or the outer protein sequence is evolved [Fig. 4**c**]. The inner–inner homotypic interactions also become stronger in both cases, although the strengthening is rather more pronounced when the inner protein is being evolved [Fig. 4**b**]. This is expected when the two coexisting condensed phases become more pure and the two components become less well mixed.^96,97^ This substantial strengthening in inner–inner interactions obtained when evolving the inner sequence indirectly results in the outer–outer interactions also becoming stronger as both phases become purer, even though the outer sequence is itself kept unchanged. By contrast, when the outer protein is evolved, the outer–outer homotypic interactions weaken even as the outer phase becomes purer [Fig. 4**a**]. Nevertheless, in both cases, we observe that the balance of interactions converges to the same behaviour across evolutionary runs: the difference in homotypic interactions becomes larger, while the heterotypic interactions become weaker and comparable in strength to the outer–outer interactions. The interaction energies in our co-evolved multiphase condensates with A1 LCD also meet the criteria we describe above [Fig. S13]. That is, multiphasicity emerges for systems with sufficiently different homotypic interactions, where the inner–inner interactions are strongest and heterotypic interactions are small but comparable to the outer–outer interactions.

Finally, we compute the interfacial free-energy densities for the liquid–vapour interface for bulk A1 LCD and its final coevolved proteins [see Methods and Fig. S14]. These results are shown in Table I and confirm the expectation that the protein with the largest surface tension with its vapour is most likely to be at the centre^46^ of the multilayered condensate. By computing the interfacial free-energy density at a range of temperatures [Fig. S14], we can extract the interfacial entropies^98^ and in turn the interfacial energies; these are also shown in Table I. The formation of the interface is energetically disfavourable; although it is in principle entropically favourable, since molecules in the liquid-like phase can gain considerable translational entropy at the interface, this contribution is relatively small. The difference in homotypic energies between species therefore also dominates the thermodynamic favourability of interface formation and determines the ordering of the layers in a multilayered condensate.

**Table I.**
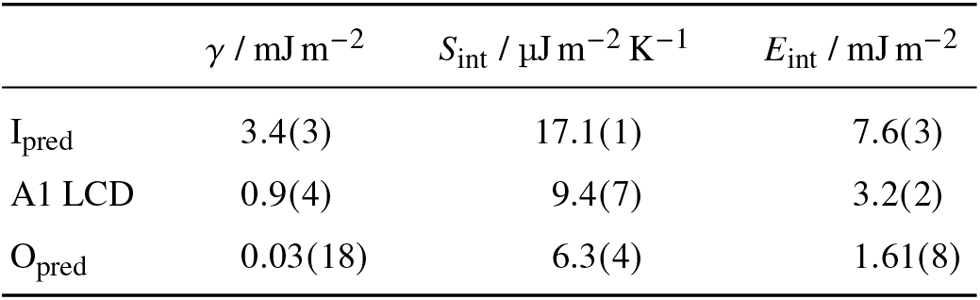
Interfacial thermodynamic parameters. Interfacial free-energy density *γ* at 250 K, interfacial entropy density *S*_int_ and interfacial energy density *E*_int_ for A1 LCD and its final coevolved proteins with maximum fitness when (i) the evolved protein is on the inside of the condensate (I_pred_) and A1 LCD is on the outside and (ii) the evolved protein is on the outside of the condensate (O_pred_) and A1 LCD is on the inside. Errors in brackets apply to the least significant digit and give the standard errors of the fitting parameters [Fig. S14].

Overall, our analyses explain why residues that increase the difference in interaction strengths between the two sequences improve multiphasicity. These results support previous studies which found that multiple immiscible phases form when there is a sufficient difference in interaction strength between the components in the two phases.^35,36,45,48,49,94^

### D. Sequence patterning is only sometimes important for multiphasicity

The patterning of interacting amino-acid groups plays an important role in determining the phase behaviour of intrinsically disordered proteins (IDPs).^70,99^ For example, the range of stability of A1 LCD condensates was shown to depend on the number and patterning of aromatic residues, which act as stickers in the ‘stickers-and-spacers’ framework.^99,100^ More uniform distributions of stickers were found to promote phase separation of A1 LCD and to decrease the propensity to form aggregates.^99^ However, in our initial evolution runs and coevolution runs with A1 LCD, we have shown that the final evolved sequences in independent repeats of the same run, despite having similar overall compositions, can differ considerably in terms of the patterning of the more strongly interacting sticker residues.

To investigate the importance of the patterning of different residues in determining the degree of multiphasicity of these two-component systems, we shuffle the final evolved sequence with maximum fitness by rearranging residues of interest (e.g. stickers or spacers) whilst keeping the overall composition of the sequence unchanged, and compute the density profiles and fitness of shuffled sequences to examine the effect of shuffling on phase behaviour. In our analyses, we consider as stickers all the aromatic residues (F, Y, W), the neutral residues with π electrons in the side chain (N, Q) and arginine (R), and the remaining residues as spacers.

Unexpectedly, when we shuffle (in multiple different ways) the evolved sequences stemming from runs where one sequence is evolved and the other is kept unchanged, the multiphasicity, and hence the fitness, are not notably altered [Fig. S16**a,b**]. Even in the extreme cases where the evolved sequence is sorted such that the residues are rearranged in order of increasing *ε*_*i*,Mpipi_ values, with all the strongly interacting sticker residues clustered together at one end of the protein, the multilayered structure was still maintained, albeit with a drop in fitness indicating a lower degree of multiphasicity in some cases. This would suggest that for these sequences, the patterning of the stickers and spacers has a minimal effect on the formation of the two coexisting phases, and that it is only the overall composition of the sequence that determines whether the two proteins will mix into one homogeneous phase.

However, for the coevolution runs with A1 LCD, we see patterning-dependent behaviour: a sorted sequence, with stickers at the ends of the protein molecules, results in rather different phase behaviour [Fig. 5**a**] compared to the original coevolved sequence [Fig. 2**c**]: in the sorted case, the sticker-rich ends interact so strongly that they form a locally crystalline structure. Interestingly, for the case where A1 LCD is at the centre of the multilayered condensate, shuffling just the positions of the spacers in the evolved sequence results in a lower degree of multiphasicity [Fig. 5**b**] relative to the system with the original evolved sequence [Fig. 2**b**]. The heterotypic interactions become stronger and the difference in homotypic interactions of the two sequences decreases after shuffling, consistent with the lower degree of multiphasicity observed. The fact that different spacers do not give rise to identical phase behaviour has recently been investigated by Bremer et al.,^95^ and similarly, in this case, we find that it is not just the distribution of stickers versus spacers that is important, but also both the identity of the spacers and their arrangement along the sequence.

**Figure 5.**
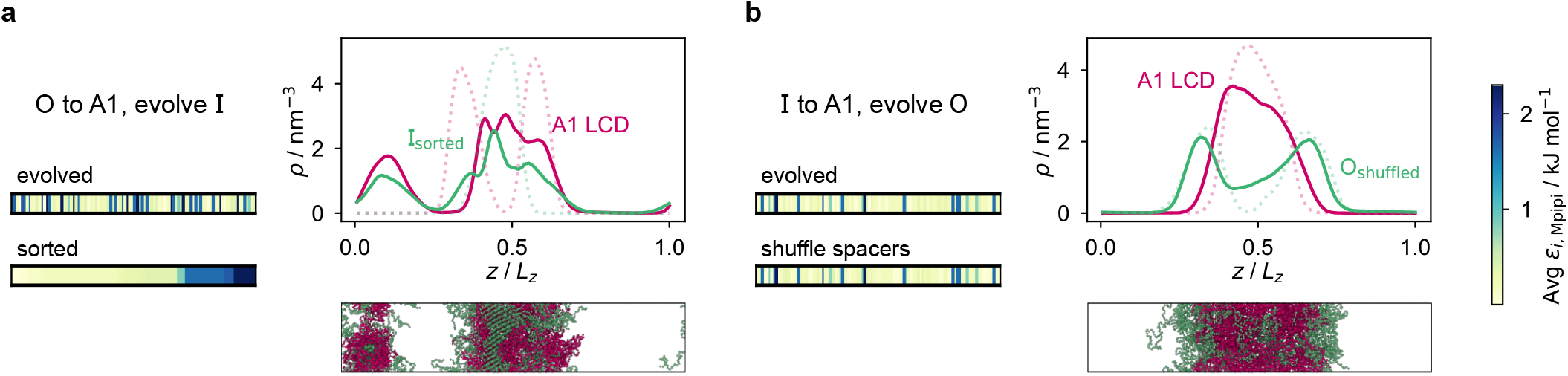
Effect of sequence patterning on the formation of multilayered condensates. **a** Density profile of the final evolved system with maximum fitness in the coevolution run where A1 LCD is designed as the outer sequence, but with the residues in the final evolved sequence I_pred_ rearranged in terms of increasing *ε*_*i*,Mpipi_. The sorted sequence with all the strongly interacting residues clustered at one end results in rather different phase behaviour. **b** Density profile of the final evolved system with maximum fitness in the coevolution run where A1 LCD is designed as the inner sequence, but with spacer residues (see text) in the final evolved sequence O_pred_ shuffled randomly. The sequence with the spacers shuffled resulted in a decrease in multiphasicity compared to that of the original evolved sequence predicted by the coevolution approach. Dotted lines represent the density profile of the original evolved system with the unshuffled partner sequence.

Although in some cases patterning of amino-acid residues does not affect the phase behaviour much, it does in others. It would be helpful to anticipate the conditions where patterning is likely to be important. Our tests suggest that if the partner protein’s sequence is repetitive with low compositional diversity, the relevant interactions can occur anywhere along the chain, reducing the need for a particular patterning of interactions to maintain phase separation. For example, for the case where we evolve the outer sequence and the inner protein is I = (FAFAA)_10_, which is highly repetitive, the multiphasicity is unaffected by shuffling or sorting the outer protein [Fig. S16**a**]; however, if we sort the inner sequence to give I = F_20_A_30_, thereby removing the repetition while maintaining the overall composition, this results in a substantial loss of multiphasicity [Fig. S17]. However, by contrast, the protein partner of the analogue where the inner sequence is evolved is not especially repetitive [see protein I_i_ of Fig. 3**b**], but patterning is nevertheless not especially important [Fig. S16**b**]. Another obvious difference between the sequences investigated is their length, and it may appear that with shorter proteins (such as those investigated in Fig. S16**a,b**), all relevant residues are spatially sufficiently close that the same interactions dominate irrespective of their precise position in the sequence. However, the behaviour of systems where A1 LCD is partnered with a 50-residue strand (cf. Fig. S7) is almost identical to the case of 100-residue strands shown in Fig. 5 and Fig. S16**c,d**, suggesting that length alone is not sufficient to rationalise the difference in behaviour.

Since it is difficult to know a priori when the patterning of residues is likely to affect the phase behaviour of multilayered condensates, the use of a genetic algorithm and the coevolution approach as a predictive tool, where any relevant patterning is optimised alongside the interaction strengths, is especially attractive.

## III. DISCUSSION

We have developed a computational approach to design multi-component multilayered condensates that contain a target protein of interest. Our approach integrates a genetic algorithm, anchored in an innovative fitness function for efficient evolution of multiphasicity, with our near-quantitative residue-resolution coarse-grained protein model, Mpipi.^83^ We demonstrate the utility of our approach in a biological context by applying it to predict different protein partners capable of forming two-component multiphase condensates when mixed with A1 LCD or its variants. We show that our method can be adapted to produce condensates that concentrate the protein of interest (e.g. A1 LCD) either at the centre of the multilayered condensate or in the outer layer, as desired.

In addition to enabling the design of multiphase condensates, our approach helps uncover the biophysical mechanisms that drive the formation of complex multilayered organisations. In all cases, we find that multiphasicity in multi-component protein systems is favoured if the difference between homotypic interactions among different components is large, and the strength of heterotypic interactions is small but comparable with that of the weaker homotypic interactions in the mixture. In a two-component system, proteins that establish stronger homotypic interactions are concentrated at the core of the multiphase condensate, as saturating their bonds enhances the overall enthalpic gain for condensate formation. Consistently, the outer layer of the multiphase condensate is formed by the proteins that establish weaker homotypic interactions, as this reduces the overall interfacial free-energy density of the two-component system. In some cases, amino-acid sequence patterning is important in controlling the phase behaviour, and shuffling the evolved sequence results in a notable decrease in the homotypic interaction strengths between each of the two components.

Since the genetic algorithm is coupled to a residue-resolution coarse-grained model for proteins, the accuracy of our predictions is contingent on that of the model. Reassuringly, we have previously demonstrated that Mpipi reproduces the experimental phase diagrams of A1 LCD and its variants with near-quantitative accuracy, achieves excellent agreement with experiments probing the phase behaviour of naturally occurring proteins (i.e. FUS, Ddx4, LAF-1 and their variants) and of polyR/polyK/polyU mixtures, and predicts experimental radii of gyration of a large set of IDPs with high accuracy.^83^ We therefore expect the predictions of our approach to be robust against experimental validation, which is an essential next step. An important factor to consider for such validation is that the specific amino-acid sequences we predict to from multiphase condensates are only applicable at the fixed temperature and salt concentration at which the simulations are run; however, these could of course be changed to design multilayered condensates that are stable under different conditions. One possible limitation of using residue-resolution coarse-grained models like Mpipi is that these are typically unable to consider the emergence of secondary or tertiary structural transitions from specific changes in amino-acid sequence. In this regard, our evolutionary approach is flexible enough to incorporate knowledge-based constraints to bypass selected patterns of amino-acid sequences known to favour, for instance, the folding of protein regions into α-helices or β-sheets in specific contexts, or limit the number of certain residues such as cysteine, which forms disulphide bridges. Our approach can also easily be modified to consider special requirements for each protein system by introducing further constraints in the algorithm; for instance to introduce tailored replacement probabilities and outcomes for different residues (e.g. to limit mutations to stickers, only allow mutations of charged to charged residues, or enforce a given pattern of aromatics) or protein regions (e.g. to avoid mutations at the N-terminus or to favour the concentration of aromatics at the centre) when proposing mutations.

While we have investigated multiphase condensates comprising only of two protein components, our evolutionary approach is transferable to multi-component systems with a larger number of components, and can also easily be extended to study the effect of RNA or post-translational modifications. In turn, our method expands the repertoire of tools available to gain molecular insight into LLPS in complex biological cellular functions. Our approach therefore presents new opportunities for designing multilayered condensates, probing more closely the underlying physicochemical factors that lead to their formation and, ultimately, deciphering the missing links to their function inside cells.

## IV. METHODS

### A. Genetic algorithm and fitness function

In our implementation of the genetic algorithm, we maintain a population of 20 sequences at each round, alongside a partner protein sequence that is not being evolved. To generate the initial population, we mutate the initial sequence by replacing, with 0.05 probability, each residue with a new one chosen from the 20 canonical amino acids with uniform probability. Each individual sequence ***x*** is assessed with a fitness function,

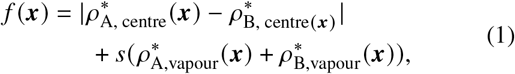

where 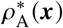 and 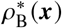 are the average dimensionless number densities of the two different protein sequences A and B in the two-component mixtures. 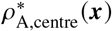 and 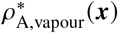 denote the number density of protein A in the core of the multilayered condensate and in the dilute phase respectively, with analogous expressions for protein B [Fig. 1**a**]. Details of how these regions are determined are discussed in Sec. S10.1. The number densities are non-dimensionalised by dividing them with an appropriate unit, e.g. *ρ*^*^ = *ρ*/nm^−3^, although the choice of unit is immaterial, since only the relative ordering in fitness is important, not the absolute numerical value. Finally, as discussed in the main text, the parameter *s* determines the trade-off between obtaining two stable liquid-like phases to give a stable multilayered condensate and the difference in compositions between the two phases. The value of *s* can be tuned as necessary depending on the specific system of interest; here, we have used *s* = 0, 0.5, 1 and 5.

Once the fitness of each individual is determined, we use the tournament selection algorithm^52,101^ to select the fittest parents to cross over. We also apply a round of random mutations to explore previously unsampled regions of sequence space. Finally, we use a weak population replacement scheme to generate the population of sequences for the next round of the genetic-algorithm run. Our genetic-algorithm implementation is detailed in full in Ref. 70.

The fitness of each individual in the population is computed when it is first encountered, e.g. following a mutation or crossover; however, when a systematic change is made to the partner sequence in coevolution runs, this too affects the phase behaviour and the fitness of all individuals in the population must therefore be recalculated. In our coevolution runs, we do this at 5-round intervals, at which we change ∼10 % of the residues of this partner protein to the target protein sequence, with residues to be changed chosen randomly from those that have not yet been changed with uniform probability.

### B. Simulation details

To simulate protein chains, we use the Mpipi residue-resolution sequence-specific coarse-grained model,^83^ combining (i) harmonic covalent bonds between residues, (ii) the Wang–Frenkel potential^102^ to account for non-bonded interactions between amino acids and (iii) Debye–Hückel electrostatic interactions.^103^ The Mpipi potential was shown to model LLPS of intrinsically disordered proteins well.^83^ For each amino-acid pair *i j*, the Mpipi model defines a Wang–Frenkel well-depth (*ε*_*i j*_), a characteristic length scale (*σ*_*i j*_) and values for *ν* and *μ* that determine the steepness of the potential well.

We use direct-coexistence simulations^104,105^ to model the vapour phase alongside the condensed phases in the same elongated tetragonal simulation box with explicit interfaces between phases. We use Lammps^106^ to run molecular-dynamics simulations with a typical time step of 10 fs and a coupling to a Langevin thermostat with a relaxation time of 10 ps.

We use 96 chains of each protein for the evolution runs with generic sequences with a box size of 11.4 nm × 11.4 nm ×56.9 nm. For coevolution runs, we use 45 chains of the protein that is changed to A1 LCD, and either 90 chains of 50 residues or 45 chains of 100 residues of the other protein, in a box of size 10.9 nm × 10.9 nm × 54.7 nm. Although finite-size effects were examined in Ref. 83 with similar-sized systems, we perform a sanity check by verifying that if we double the system size, the density profiles are consistent and the ordering of fitness values is the same [Fig. S20].

### C. Interfacial free-energy densities

We compute interfacial free-energy densities for the interface between the vapour-like phase and the pure condensed phase for the final evolved maximum-fitness sequences in coevolution runs with A1 LCD for both cases, i.e. where the coevolved protein is the inside or the outside protein. To do this, we use the Kirkwood–Buff expression^107^ to relate the interfacial free-energy density *γ* to the normal and tangential components of the pressure tensor, and the mean-value theorem to simplify the result^108^ for planar interfaces into *γ* = (*L*_*z*_/2(*P*_norm_ −*P*_tang_), where *γ* is the interfacial free-energy density, *L*_*z*_ is the length of the simulation box along which the interface occurs, *P*_norm_ = *P*_*zz*_ is normal to the interface and *P*_tang_ = *P*_*xx*_ = *P*_*yy*_ is the tangential pressure, and the division by 2 accounts for the fact that there are two interfaces in our simulation set-up.^109^ Although the pressure tensor has many possible definitions, from virial to mechanical expressions, the interfacial-free energy density is independent of this arbitrary choice;^110^ we use the atomic virial pressure tensor in our calculation, which gives the same results as the molecular virial.^111^

We compute only the interfacial free-energy density for the interface between the dense and dilute phases (i.e. a surface tension using our coarse-grained model), since as long as a multilayered condensate forms, the interface between the two condensed phases of different compositions is always present and therefore does not affect the thermodynamics. We assume for simplicity that the resulting phases are pure and, in this back-of-the-envelope calculation, we do not consider the possible dependence of *γ* on the interface width or the curvature of the droplet. We determine the interfacial free-energy density for each system at several temperatures. The resulting data are well fitted by a linear function [Fig. S14]; since (*∂γ ∂T* = *S*_int_), the interfacial entropy,^98^ this approach allows us to extract the interfacial energy and interfacial entropy for each component, as discussed in the main text.

## V. DATA AVAILABILITY

All relevant data are within the manuscript, its Supporting Information files and the Figshare data repository at TBD.

## VI. CODE AVAILABILITY

Lammps input scripts are available in the Figshare data repository at TBD.

## VII. ACKNOWLEDGEMENTS

We acknowledge funding from the University of Cambridge Ernest Oppenheimer Fund [PYC], the Winton Programme for the Physics of Sustainability [PYC, RC-G], the European Research Council under the European Union’s Horizon 2020 research and innovation programme [grant 803326; RC-G]. JAJ is a Junior Research Fellow at King’s College. This work was performed using resources provided by the Cambridge Tier-2 system operated by the University of Cambridge Research Computing Service funded by EPSRC Tier-2 capital grant EP/P020259/1 [RC-G, JAJ, AR]. The funders had no role in study design, data collection and analysis, decision to publish or preparation of the manuscript.

## VIII. AUTHOR CONTRIBUTIONS STATEMENT

PYC, JAJ, RC-G and AR designed the research. PYC performed the research. PYC, JAJ, RC-G and AR analysed the results and wrote the paper.

## IX. COMPETING INTERESTS

The authors declare no competing interests.

## S10 SUPPLEMENTARY INFORMATION

### S10.1 Determining the centre of the density profile

To calculate the density profiles, the simulation box is divided into 150 bins along the elongated axis, and the average density of each species is calculated within each bin. The ‘centre’ region of the initial reference system is taken as the region between where the density profiles of the two protein species intersect. The ‘vapour’ region is then defined as the 50 bins in total where the first and last bin are equidistant from the middle of the ‘centre’ region. Once these regions are quantified for the initial reference system, we fix the centre of mass of the condensate in our simulations and keep these regions constant throughout the entire genetic algorithm run.

### S10.2 Effect of weighting parameter *s* in the fitness function

**Figure S6.**
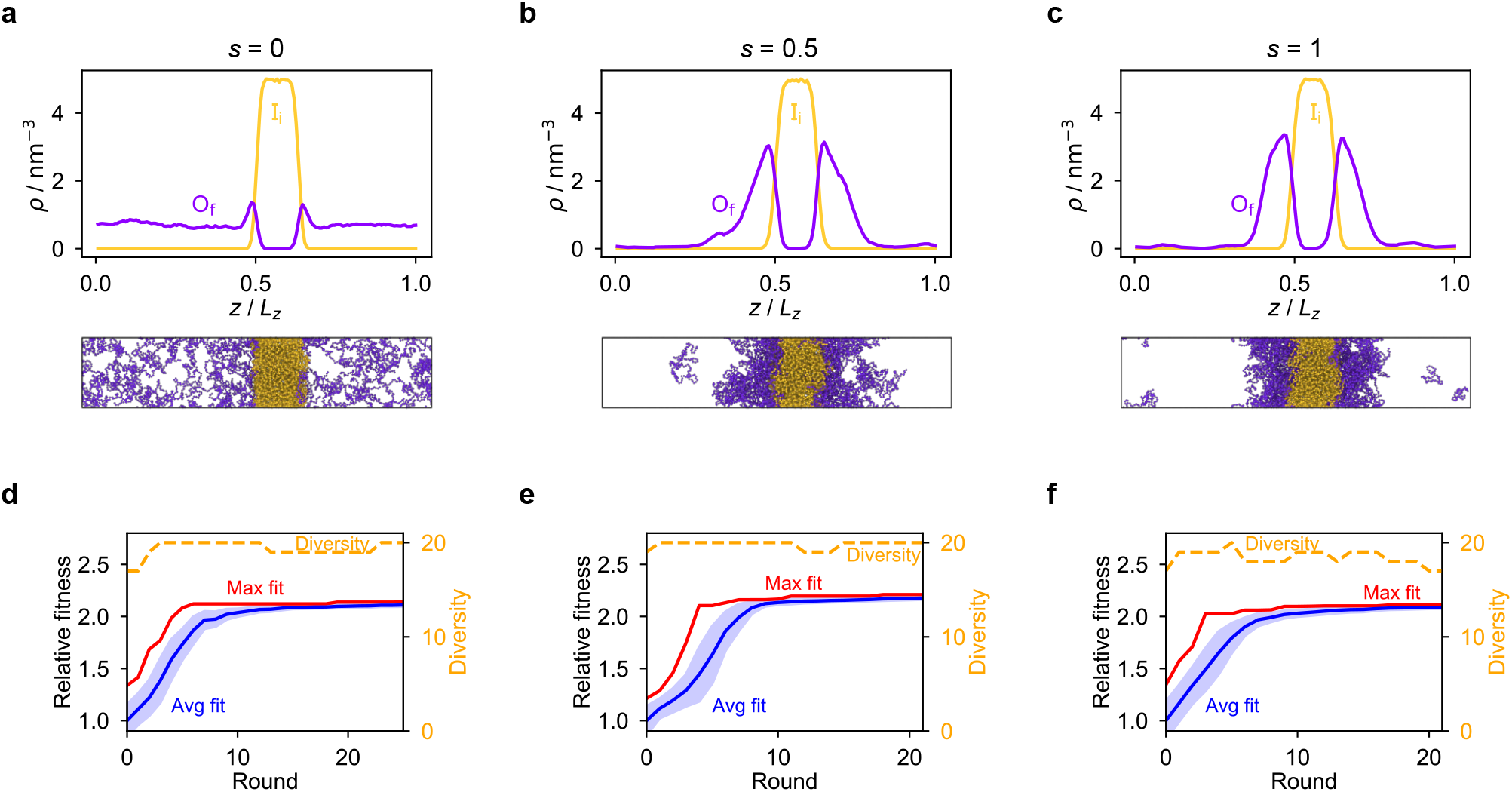
Effect of weighting parameter *s* in the fitness function. Density profiles of the final evolved systems with maximum fitness using **a** *s* = 0, **b** *s* = 0.5 and **c** *s* = 1 in the fitness function of the genetic algorithm for the evolution run where the outer sequence is evolved and the inner sequence is kept unchanged. In this case, the quality of the final result is dependent on the value of *s*. When *s* = 0, the fittest individual corresponds to a sequence which results in a system with a dense phase of one component and a dilute phase of the other, instead of two liquid-like phases with different compositions in a multilayered arrangement. To obtain the latter, *s* > 0 was needed in the evolution run. **d–f** Genetic algorithm progressions for the three cases with different values of *s*. Shaded area for the average fitness corresponds to the standard deviation across all the sequences present in the population at each round.

### S10.3 Coevolution of A1 LCD with other reference systems

**Figure S7.**
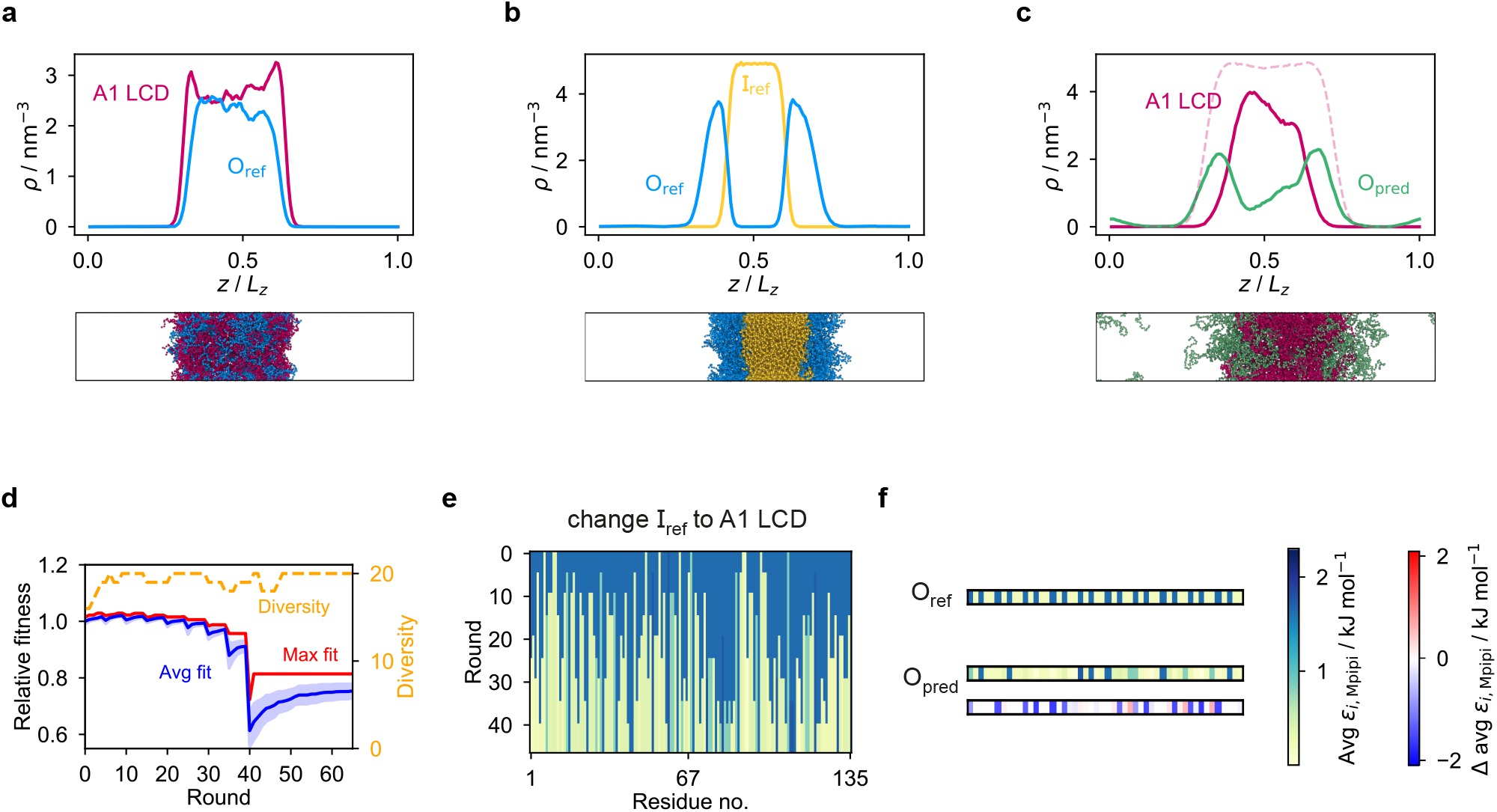
Coevolution of A1 LCD with a shorter partner protein. Coevolution run with A1 LCD designed to be the inner sequence with a shorter partner sequence of 50 residues as the outer sequence. **a** System with low multiphasicity obtained when the systematic change to A1 LCD is made directly all at once from **b** the initial reference system with high multiphasicity. I_ref_ = F_135_ and O_ref_ = FAFAA _10_. **c** Final evolved system with maximum fitness obtained from the coevolution approach. In this case, using a shorter partner sequence on the outside gives a result with a lower degree of multiphasicity compared to when a partner sequence of twice the length is used. **d** Genetic algorithm progression of this coevolution run. Shaded area for the average fitness corresponds to the standard deviation across all the sequences present in the population at each round. With a shorter partner sequence, the maximum fitness does not improve beyond round 40 after all the systematic changes has been made. **e** Illustration of how the systematic change from I_ref_ to A1 LCD is made. **f** Comparison of the final evolved sequence with maximum fitness, O_pred_, with the initial reference sequence O_ref_. To illustrate the sequences in **e–f**, we plot for each residue *i* along the sequence the absolute value and the change in the value of *ε*_*i*,Mpipi_ compared to the initial reference sequence.

**Figure S8.**
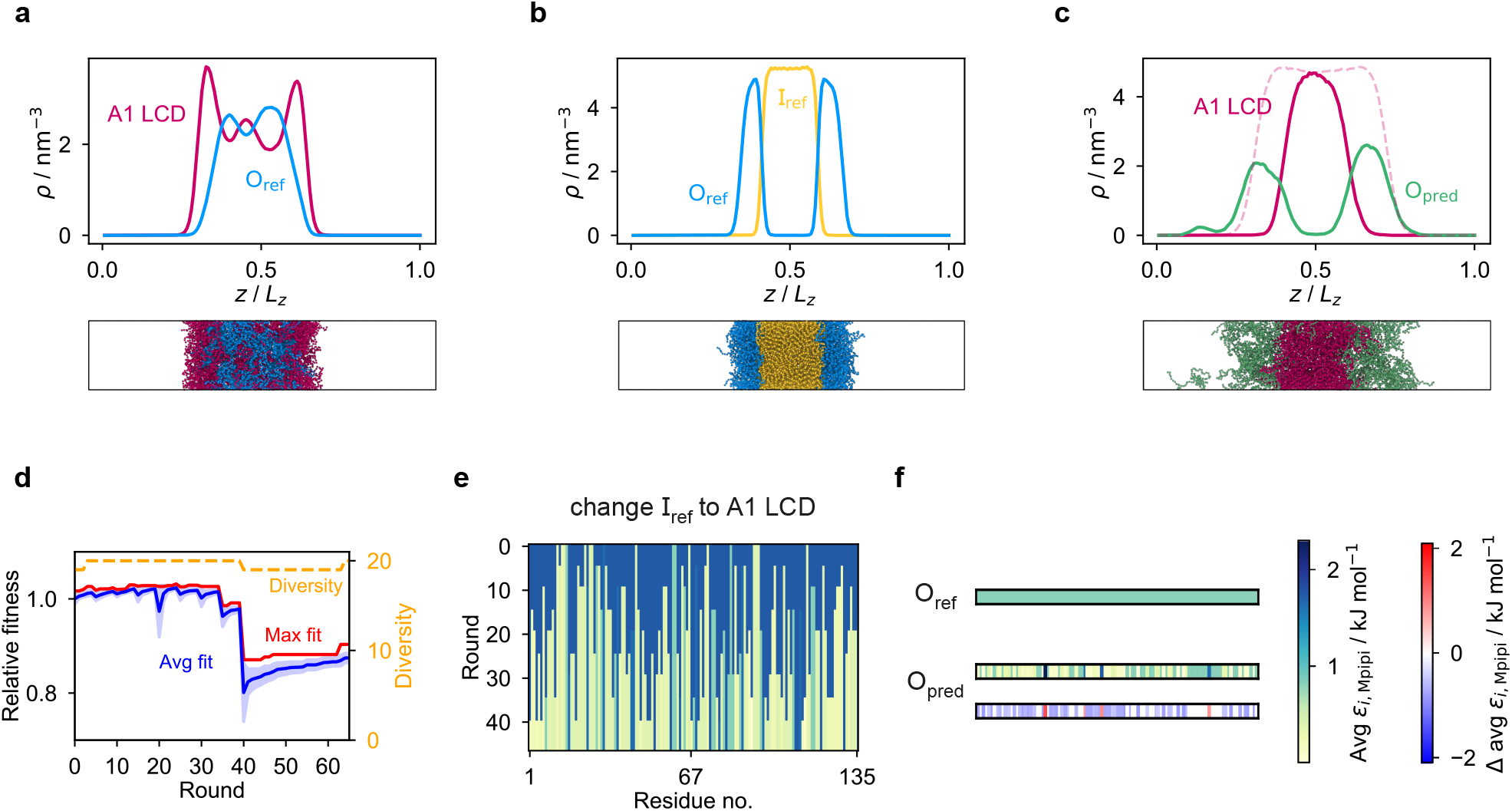
Coevolution of A1 LCD from different reference proteins. The analogue of Fig. S7 starting from a different initial reference system, namely I_ref_ = Y_135_ and O_ref_ = N_100_.

### S10.4 Systematic changes during coevolution runs

**Figure S9.**
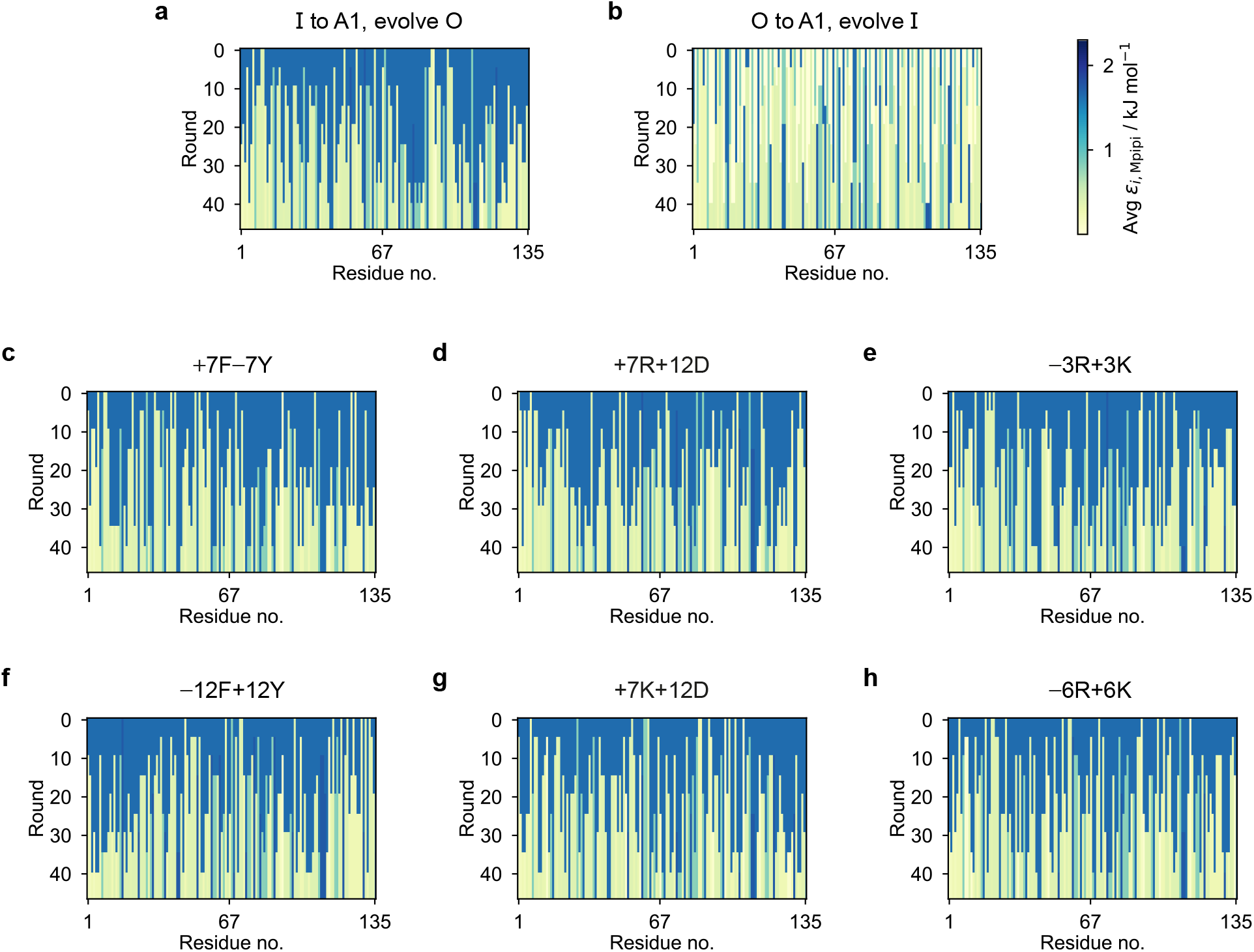
Manner in which the systematic changes are made in the coevolution runs with A1 LCD and its variants. We illustrate the sequence of the protein that is systematically changed to the A1 variant in each round by plotting the value of *ε*_*i*,Mpipi_ for each residue *i* along the sequence.

### S10.5 Genetic-algorithm progressions for coevolution runs

**Figure S10.**
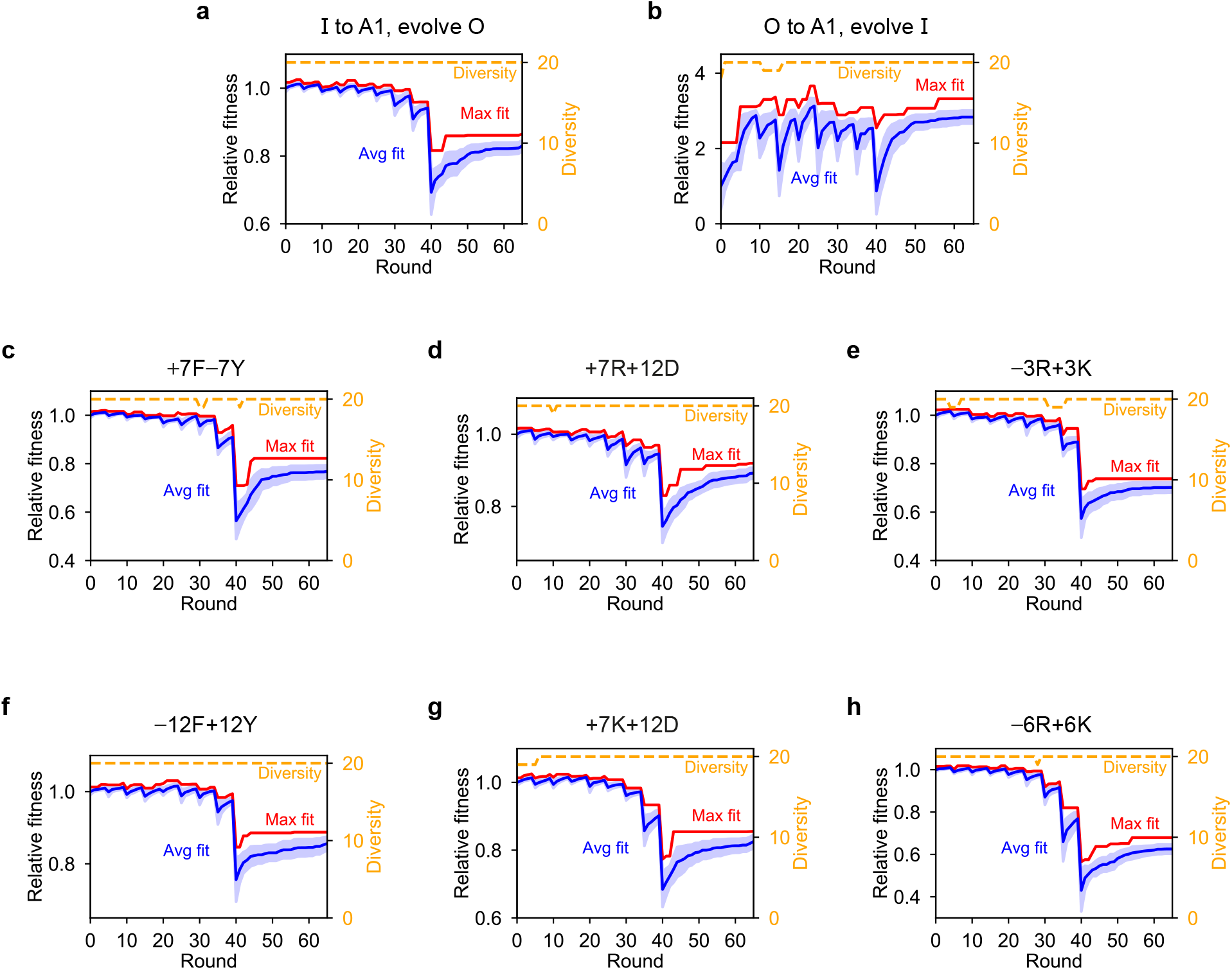
Genetic-algorithm progressions for coevolution runs with A1 LCD and its variants. Shaded areas for the average fitness correspond to the standard deviation across all the sequences present in the population at each round.

### S10.6 Coevolving A1 LCD at the centre or on the outside

**Figure S11.**
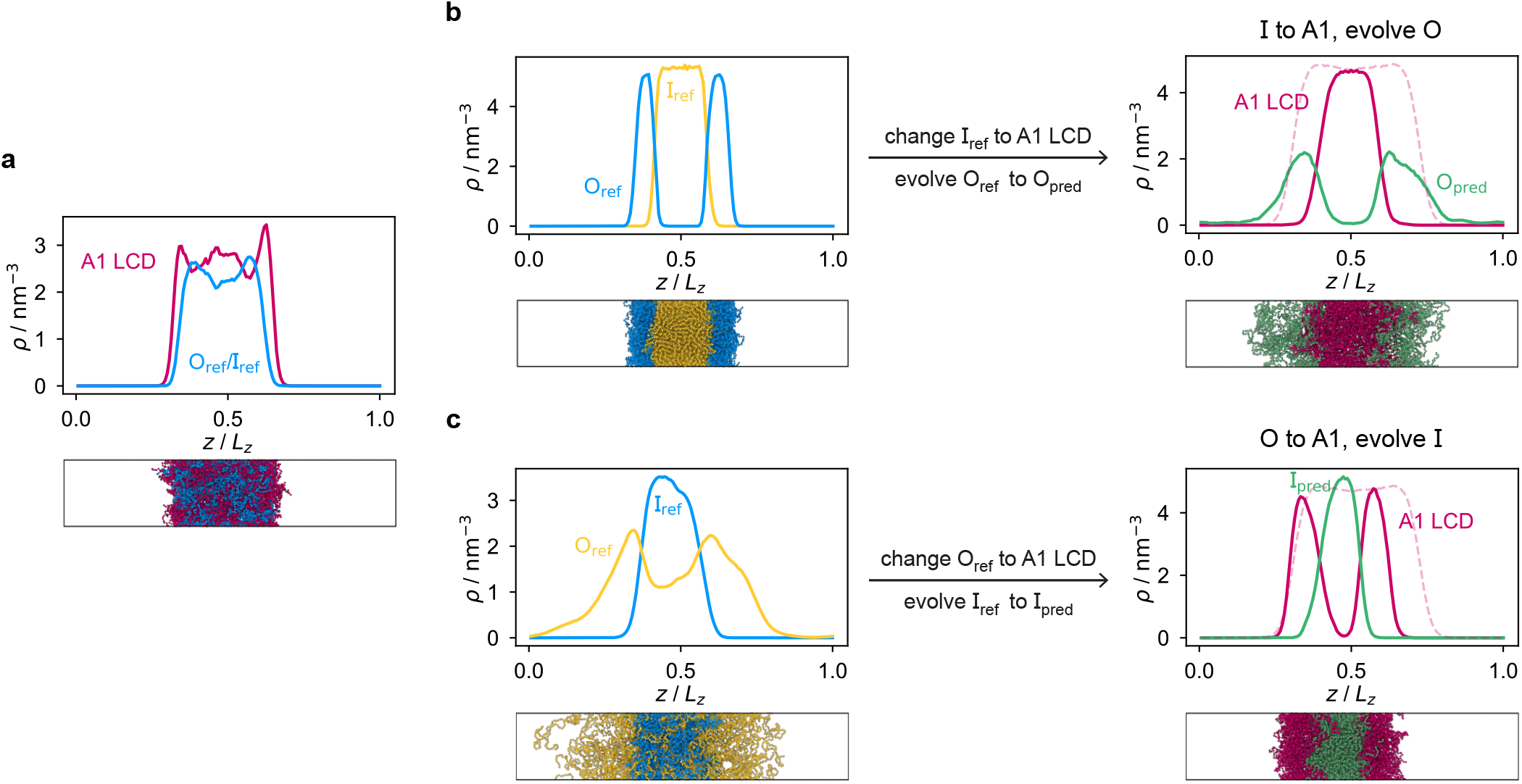
Designing multilayered condensates with A1 LCD concentrated at the centre or in the outer layer. **a** System with low multiphasicity obtained when the systematic change to A1 LCD is made directly all at once from the two initial reference systems with high multiphasicity. O_ref/_ I_ref_ = (FAFAA) _20_. **b** Density profiles of the initial reference system and final evolved system for the case where A1 LCD is designed to be the inner sequence. I_ref_ = F_135_ is systematically changed to A1 LCD and O_ref_ = FAFAA _20_ is evolved using the genetic algorithm. **c** Density profiles of the initial reference system and final evolved system for the case where A1 LCD is designed to be the outer sequence. O_ref_ = FIQII _27_ is systematically changed to A1 LCD and I_ref_ = FAFAA _20_ is evolved. The pink dashed lines in the final evolved systems of both cases is the density profile of a single-component system of A1 LCD equilibrated at the same temperature. Note that O_ref_ in panel **b** is the same as I_ref_ in panel **c**.

### S10.7 Changes in interaction energies during genetic-algorithm runs

**Figure S12.**
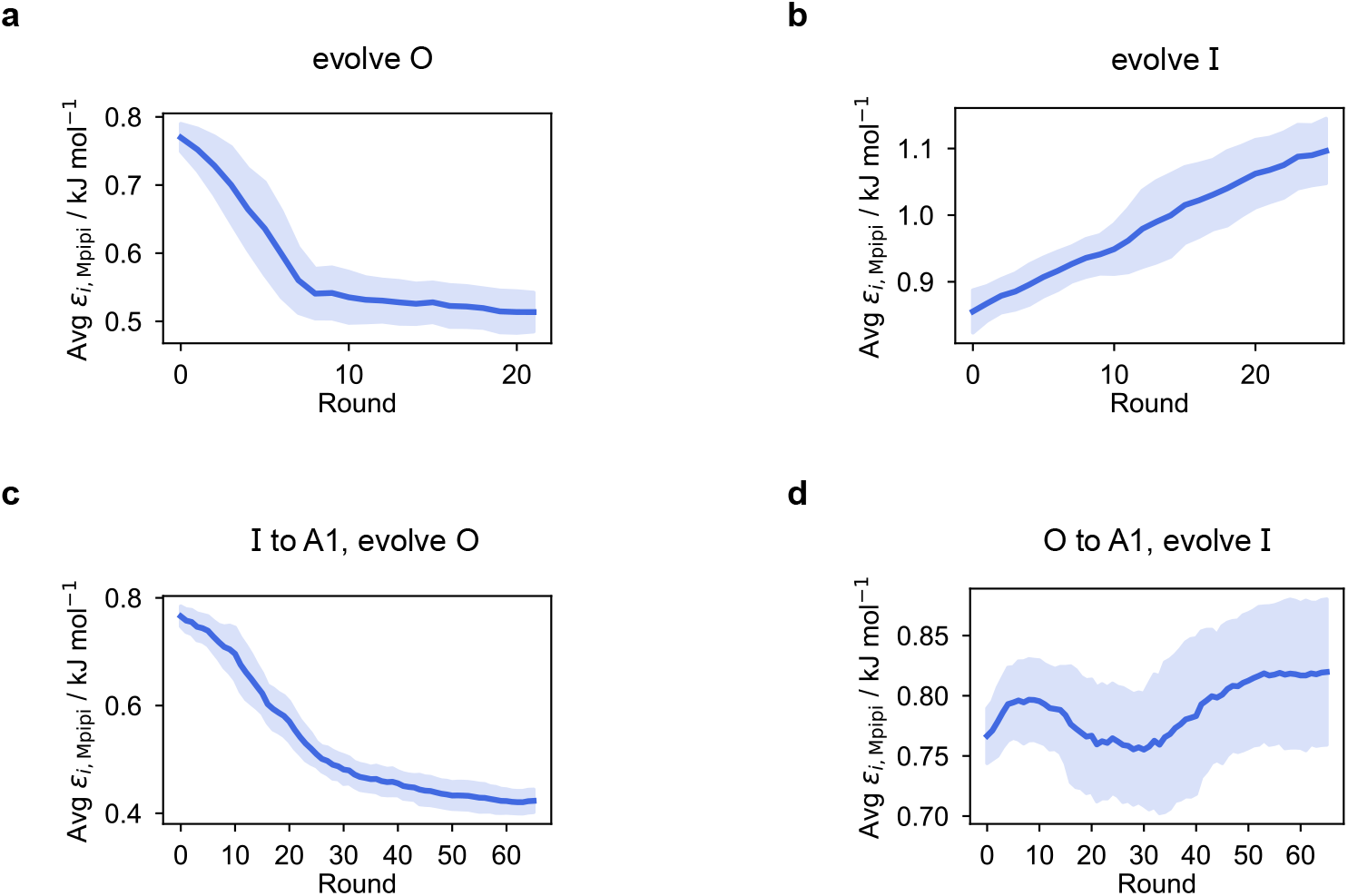
Changes in the average value of *ε*_*i*,Mpipi_. These are shown for the evolution runs where **a** the outer sequence is evolved and the inner sequence is kept unchanged and **b** the inner sequence is evolved and the outer sequence is kept unchanged, and for the coevolution runs where A1 LCD is designed to be **c** the inner sequence or **d** the outer sequence in the final multilayered system. The value of *ε*_*i*,Mpipi_ plotted is averaged over all the residues of the evolved sequences in the entire populations in three independent runs for each case. Shaded areas correspond to the standard deviation.

**Figure S13.**
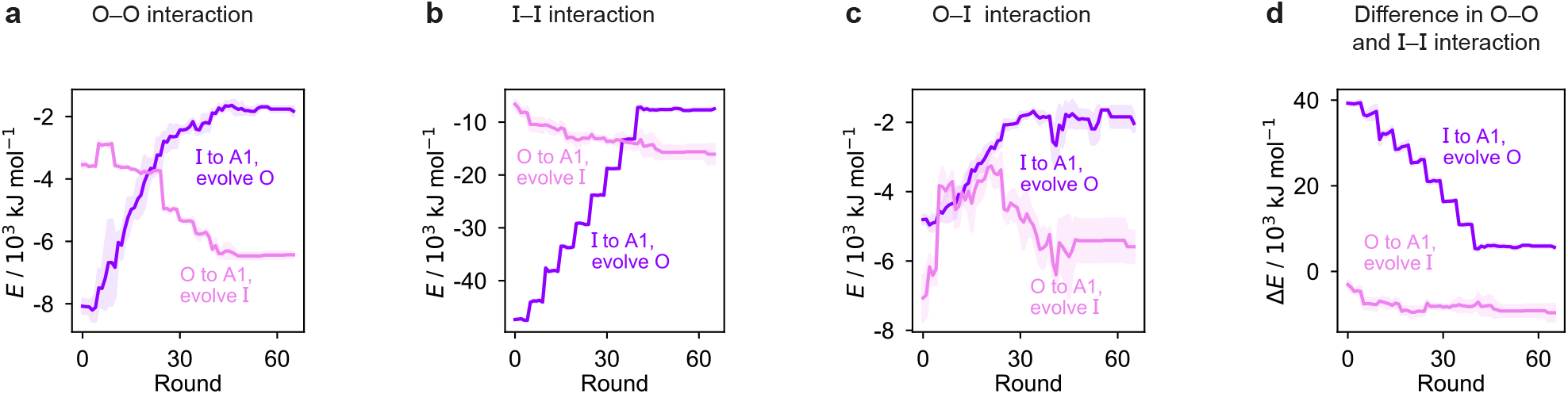
Change in the interaction energies during genetic-algorithm progression. We show these changes for intermolecular self-interactions between proteins enriched in **a** the outer and **b** the inner layer, **c** cross-interactions between the two proteins and **d** the difference in the O–O and I–I interaction energies for the system with the fittest individual in the coevolution runs where one sequence is changed to A1 LCD and the other is coevolved. Shaded areas correspond to the standard deviation across three independent runs. Although the difference in O–O and I–I interactions decreases overall, this is because of the systematic changes made to the sequence. The trends for the genetic-algorithm rounds in between those at which systematic changes occur are largely consistent with the results shown in Fig. 4.

### S10.8 Interfacial free-energy densities

**Figure S14.**
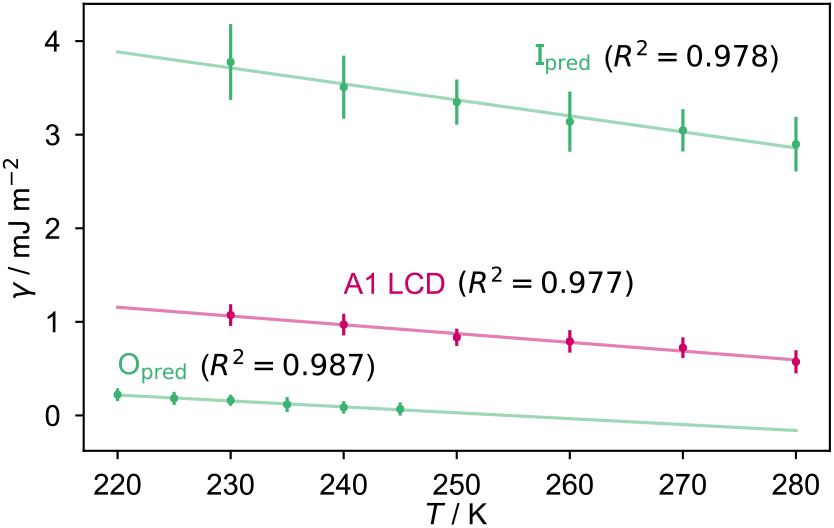
Interfacial free-energy densities as a function of temperature. Error bars correspond to the standard deviation of the interfacial free-energy density computed in several independent sections of the simulation. We use a linear fit to the data points to extract the interfacial entropy and energy for each sequence, as discussed in the main text.

### S10.9 Composition and patterning in coevolution runs

**Figure S15.**
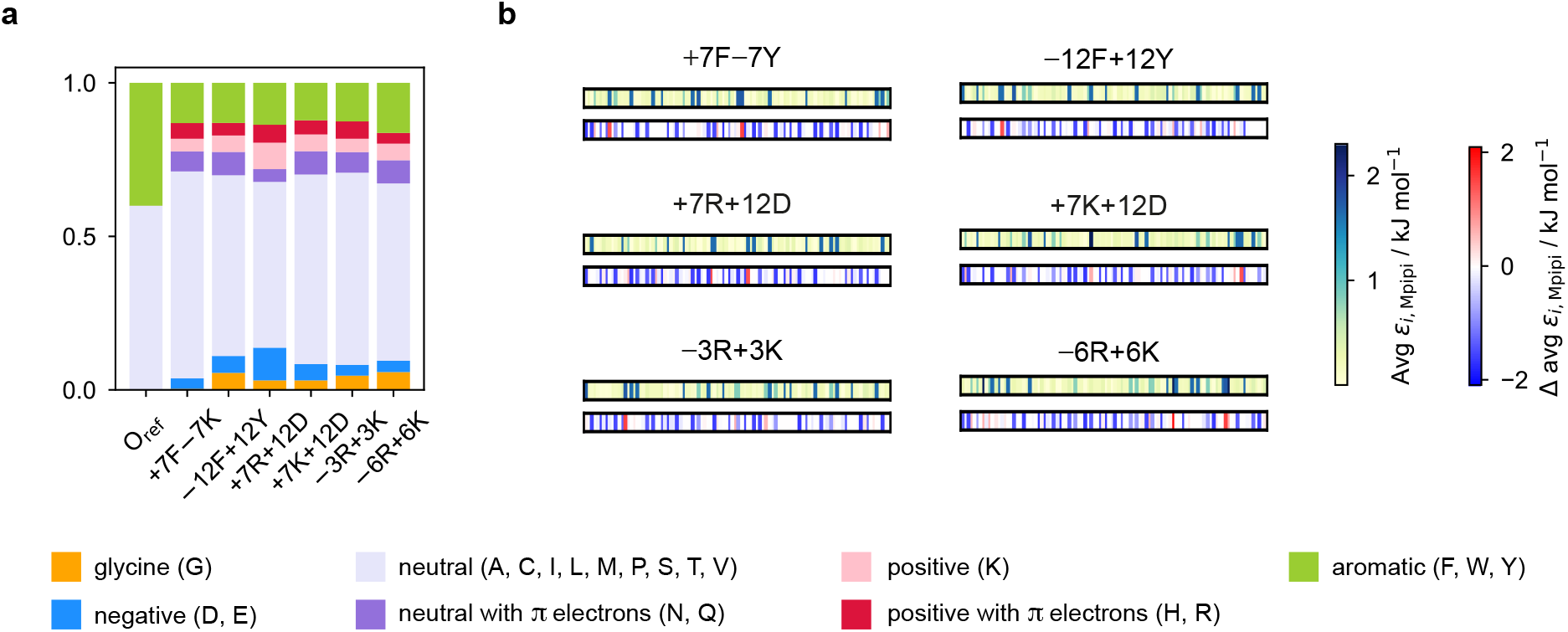
Composition and patterning in coevolution runs. **a** The amino-acid composition and **b** patterning of the final evolved sequences with maximum fitness in the coevolution runs with 6 different variants of A1 LCD designed to be in the centre of the final multilayered condensate. In **b** we plot the absolute value and the change in the value of *ε*_*i*,Mpipi_ compared to the initial reference sequence for each residue *i* along the final evolved sequence. In all cases we observe a decrease in the proportion of aromatic residues, and the final sequences contain less strongly interacting residues overall.

### S10.10 Shuffling amino-acid sequences

**Figure S16.**
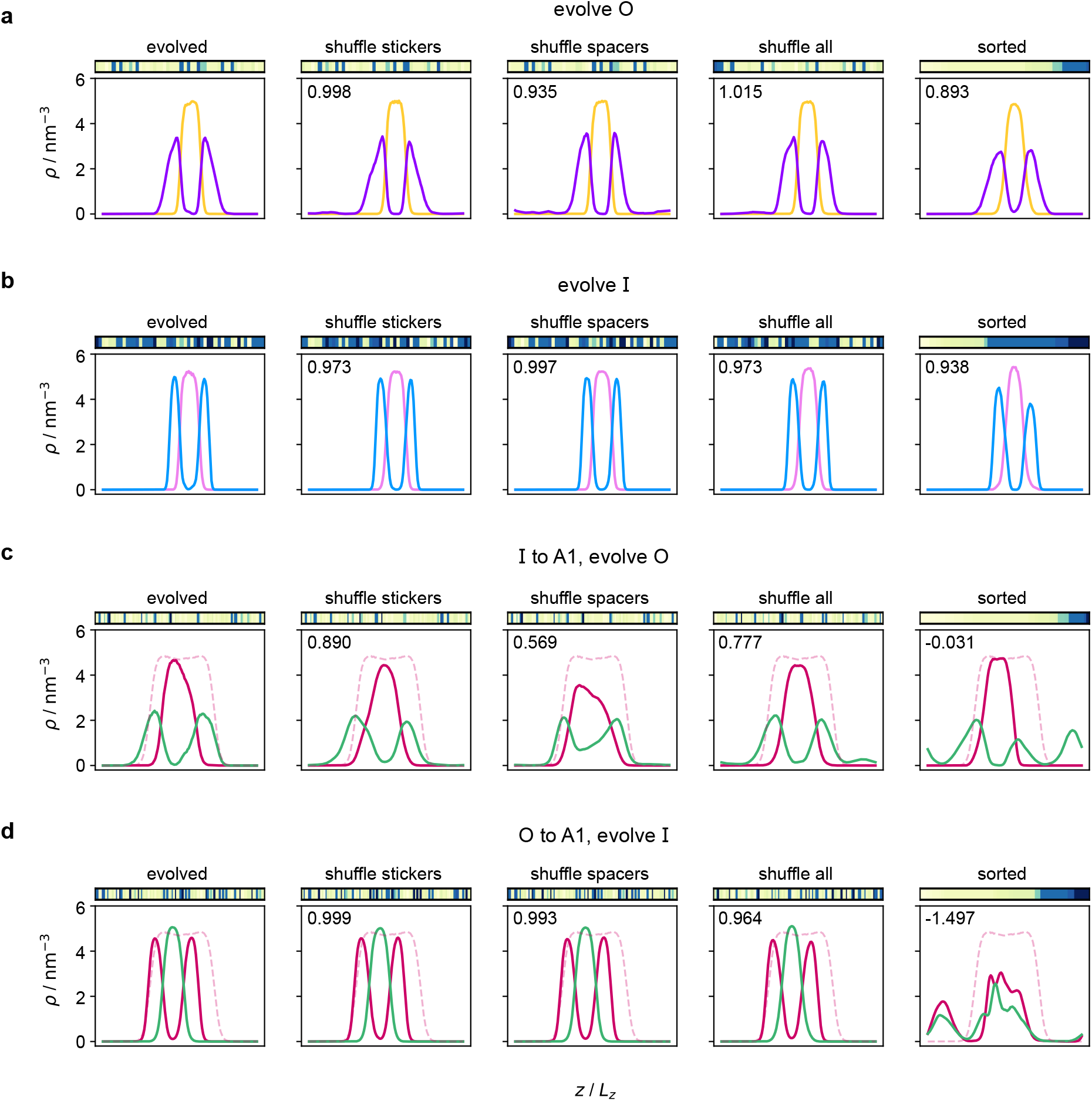
Sequence patterning only sometimes affects multiphasicity. Density profile of the final evolved system with maximum fitness in **a–b** evolution and **c–d** coevolution runs with A1 as the partner sequence, with inner or outer proteins evolved as indicated. The first column corresponds to the final evolved system with maximum fitness in each case, reproduced from the main text [**a** = Fig. 1**c**; **b** = Fig. 1**d**; **c** = Fig. 2**b**; and **d** = Fig. 2**c**]. The remaining columns correspond to different shuffles of the same sequence, as indicated, using the same notation and colours as in the main text. Just above each density profile, we show a map of *ε*_*i*,Mpipi_ values for the sequence in question, as in Fig. 5. In the top left-hand corner of each density plot, we give the fitness value relative to the evolved sequence of the left-most column. In the case of panels **a** and **b**, even if amino acids are completely sorted by their interaction strength in the sequence, the multiphasicity is not appreciably reduced, suggesting that patterning is not crucial in these cases. By contrast, even just shuffling the spacers in panel **c** leads to a considerable reduction in fitness, whilst sorting the amino acids results in very different phase behaviour in panels **c** and **d**, indicating that patterning is important in these cases.

**Figure S17.**
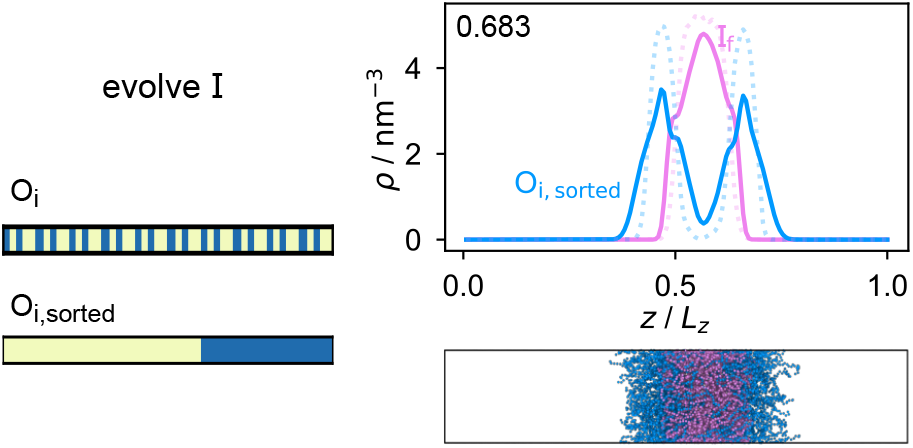
Density profile for sorted partner sequence. Density profile and simulation snapshot of the final evolved system with maximum fitness in the evolution run where the inner sequence is evolved, but with the originally fixed partner sequence O_i_ = FAFAA _10_ sorted such that residues of the same type are all clustered together. The density profile of the original evolved system is shown by the dotted lines. The number in the top left corner of the density plot is the fitness value relative to the original system with the unsorted partner sequence.

### S10.11 Testing the coevolution approach with generic sequences

As a proof of principle, we first tested the coevolution approach on simple two-component systems with generic sequences before applying it to biologically relevant proteins like A1 LCD as detailed in the main text.

For these coevolution runs, the initial reference system we used is a mixture of F_50_ and (FAFAA)_10_, which form a multilayered condensate with two compositionally distinct liquid-like phases of high purity: F_50_ concentrates in the centre while (FAFAA)_10_ remains in the outer layer. We selected two target sequences into which either one of F_50_ or (FAFAA)_10_ is systematically changed whilst the other is simultaneously evolved using the genetic algorithm. These target sequences are selected such that making the systematic change directly in one go would result in complete mixing to give one homogeneous liquid-like phase. The systematic change from the initial reference sequence to the target sequence is made by changing 10 % of the residues every 5 rounds [Fig. S18**e** and Fig. S19**e**].

In Fig. S18**b–c** and Fig. S19**b–c**, we show the density profiles and snapshots at the start and end of the coevolution runs; the final systems clearly exhibit two liquid-like phases of different composition, demonstrating that the coevolution approach is able to predict a partner sequence that can form a multilayered condensate with another sequence of interest. The approach is successful both when the target sequence is designed to be concentrated at the centre or on the outside of the multilayered condensate. The genetic algorithm progressions in the two cases are shown in Fig. S18**d** and Fig. S19**d**. The rounds where a drop in the average and maximum fitness is observed broadly corresponds to rounds where systematic changes are made.

The final evolved sequence with maximum fitness in the two different evolution cases in terms of the approximate interaction strengths of the residues are shown in Fig. S18**f** and Fig. S19**f**.

The behaviour of these generic sequences across the genetic-algorithm progression both in terms of fitness and in terms of the types of interaction that favour multiphasicity is very similar to the case of coevolution with A1 LCD discussed in the main text, even though the generic sequences we have used are considerably shorter than A1 LCD, suggesting that these results are largely independent of the specifics of the amino-acid sequences in question.

**Figure S18.**
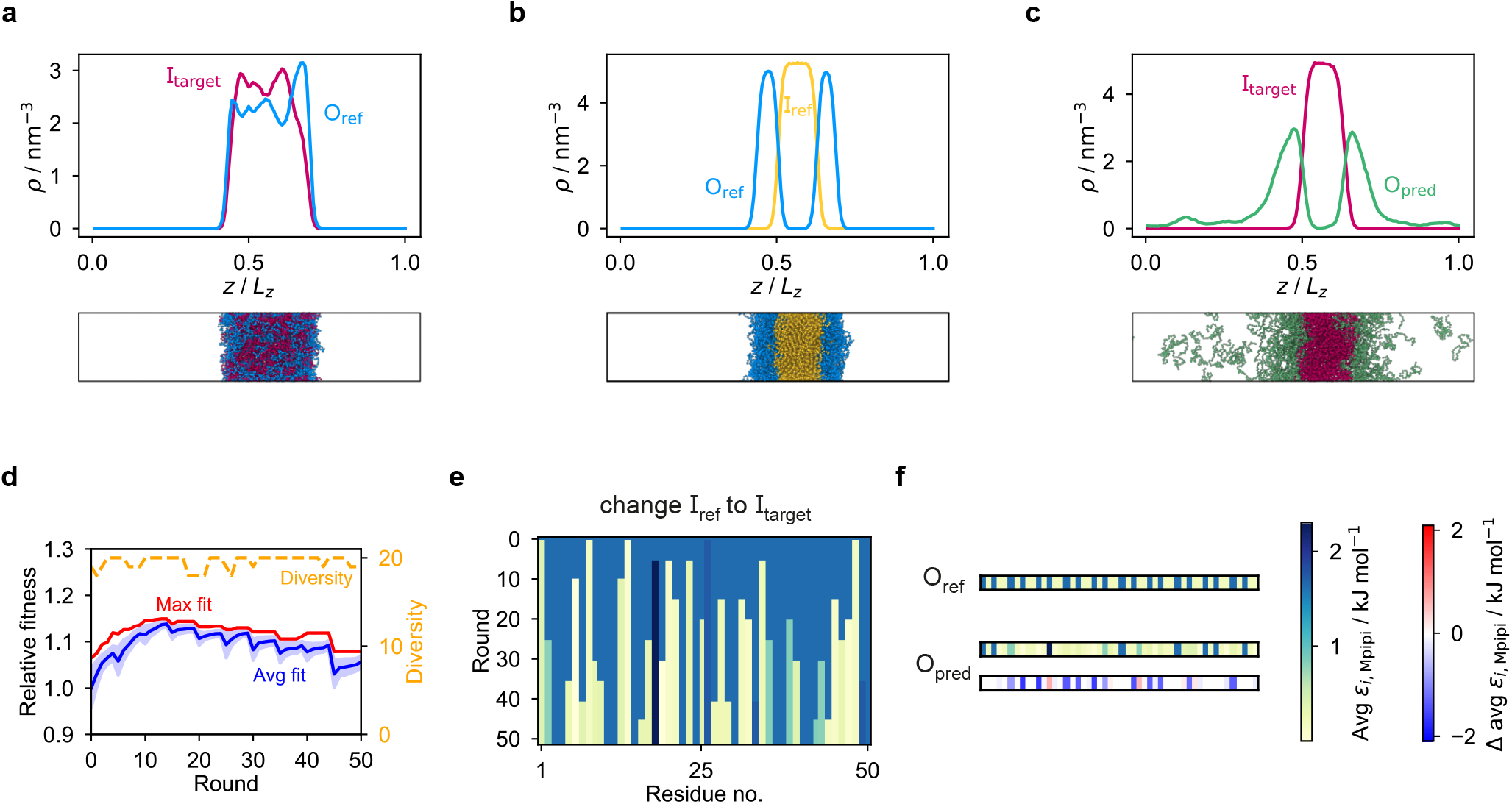
Coevolution with generic sequences (I). Coevolution run with generic sequences where the inner sequence is systematically changed and the outer sequence is evolved. I_ref_ = F_50_, O_ref_ = FAFAA _10_ and I_target_ = mutated FAFAA _10_, with a mutation probability of 0.60. **a** System with low multiphasicity obtained when the systematic change is made directly all at once from **b** the initial reference system with high multiphasicity. **c** Final evolved system with maximum fitness. **d** Genetic algorithm progression of this coevolution run. Shaded area for the average fitness corresponds to the standard deviation across the whole population at each round. **e** Illustration of how the systematic change from I_ref_ to I_target_ is made. **f** Comparison of the final evolved sequence with maximum fitness, O_pred_, with the initial reference sequence O_ref_. To illustrate the sequences in **e–f**, we plot the absolute value and the change in the value of *ε*_*i*,Mpipi_ compared to the initial reference sequence for each residue *i* along the sequence.

**Figure S19.**
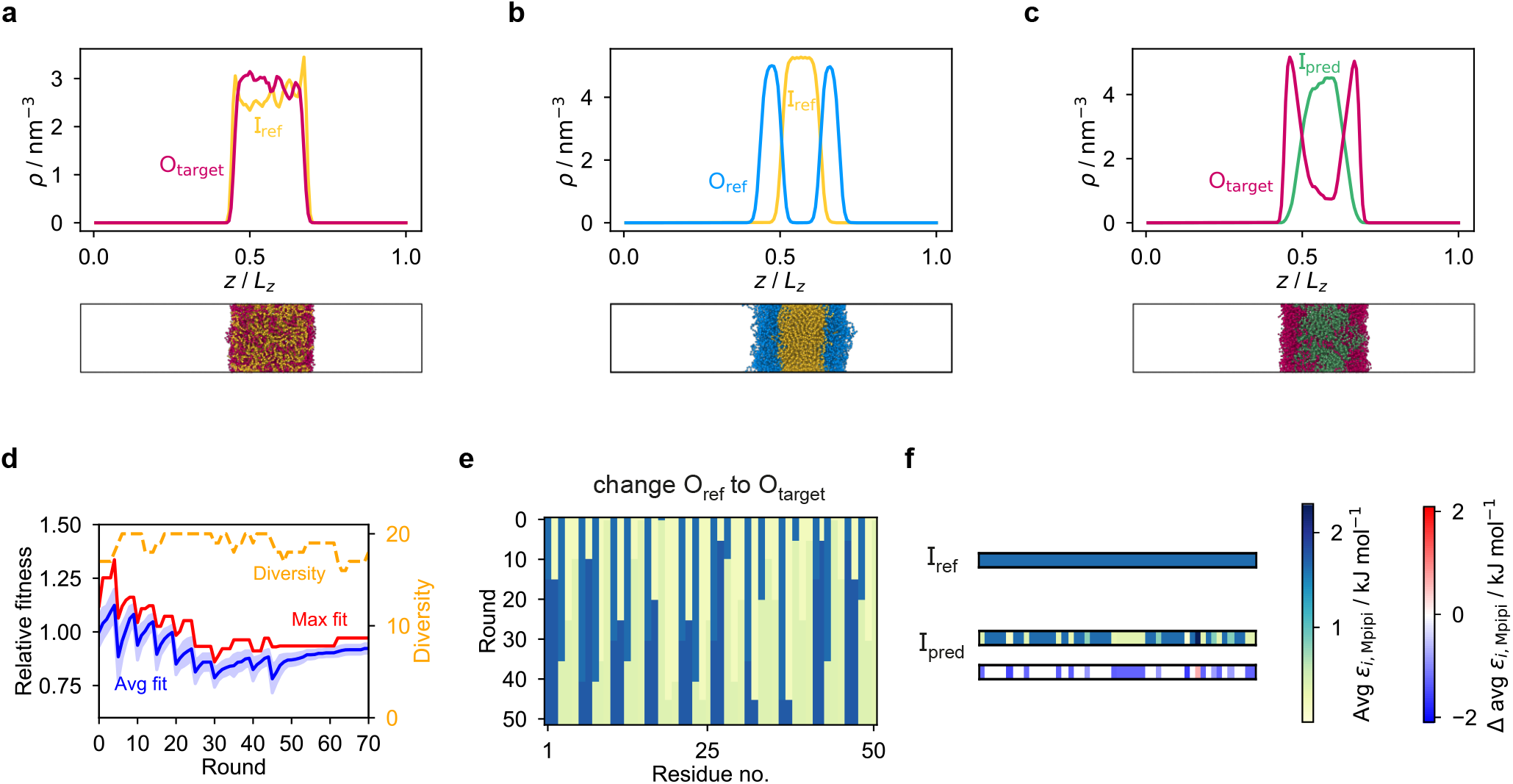
Coevolution with generic sequences (II). Analogue of Fig. S18 for a coevolution run with generic sequences where the outer sequence is systematically changed and the inner sequence is evolved. I_ref_ = F_50_, O_ref_ = (FAFAA)_10_ and O_target_ = (YYGGR)_10_.

### S10.12 Finite-size scaling

**Figure S20.**
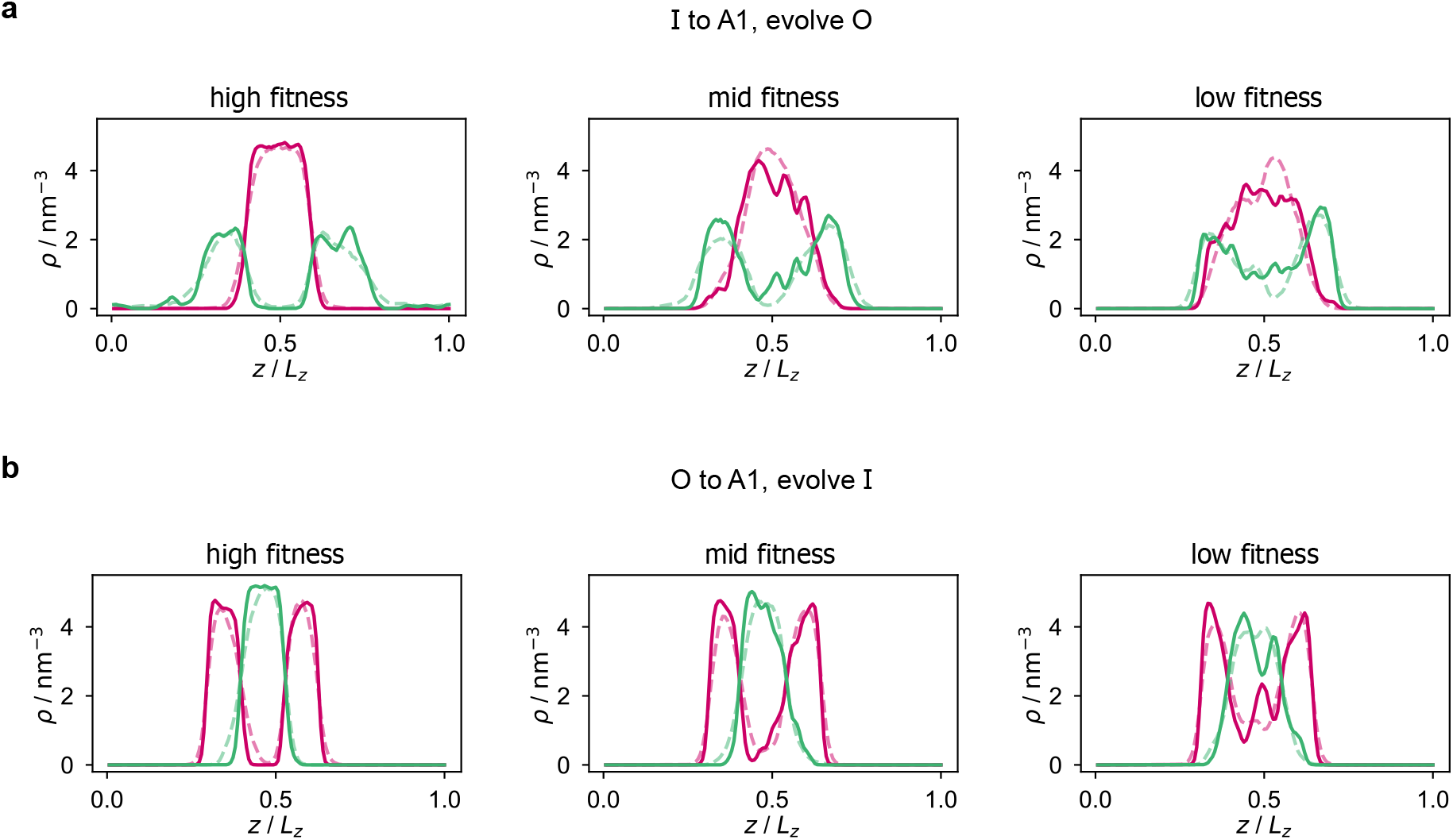
Finite-size scaling. We have computed density profiles for systems obtained in coevolution runs with A1 [cf. Fig. 2] at twice the size of the systems reported in the main text, i.e. with 90 molecules of A1 LCD, 90 chains of 100 residues each of the partner protein in a box of size 10.9 nm × 10.9 nm × 109.4 nm. The resulting density profiles, using the same colours and notation as in Fig. 2], are within typical noise of the analogues for the original system size (shown with dashed lines, which are scaled by a factor of two along the horizontal direction to ease comparison). The behaviour observed is similar irrespective of how fit the systems are, and the fitness ordering is maintained when the system size is doubled, suggesting that finite-size effects are unlikely to dominate the phase behaviour.

### S10.13 Listings of protein sequences

We list below the sequences of the proteins from the figures reported of the main text. For the final evolved proteins, we highlight changes from the initial sequence in red. The sequences given here correspond to those with the maximum fitness only. In the supporting data, we provide listings of all sequences in the population of the final round of the genetic-algorithm runs.

#### S10.13.1 Figure 1

The protein O_i_ is (FAFAA)_5_. The protein I_i_ is F_50_ with added random mutations with a probability of 0.6.

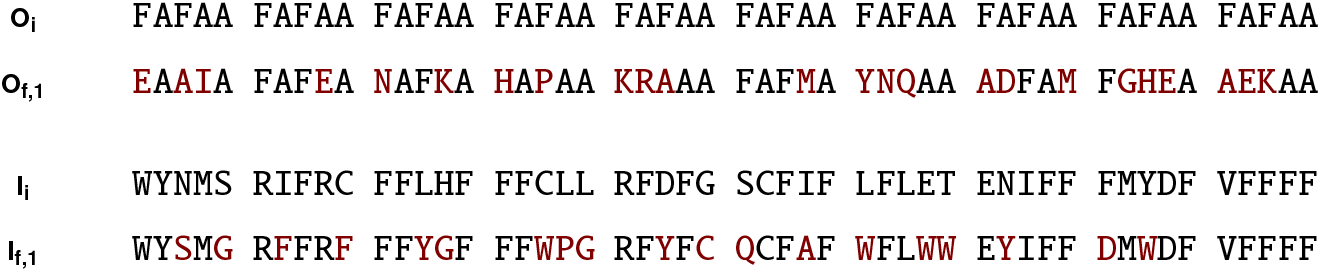

#### S10.13.2 Figure 2

We list below the sequences of the proteins shown in Fig. 2 of the main text.

We first list the sequence of A1 LCD and its variants, using the nomenclature of Bremer and co-workers.^95^ The amino-acid residues different from the wild type are highlighted in blue.

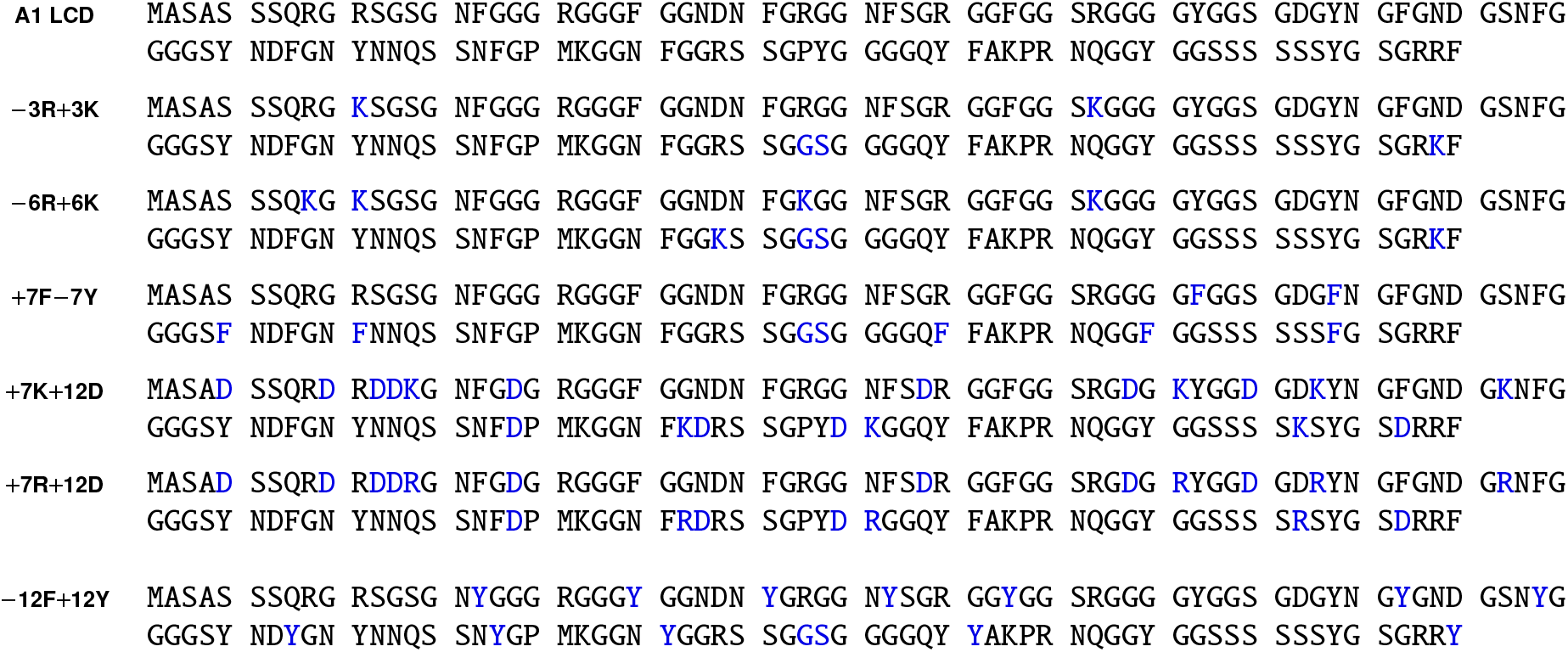

*Evolution of the outer protein when A1 LCD is at the centre.* In this case, I = F_135_ and O = (FAFAA)_20_.

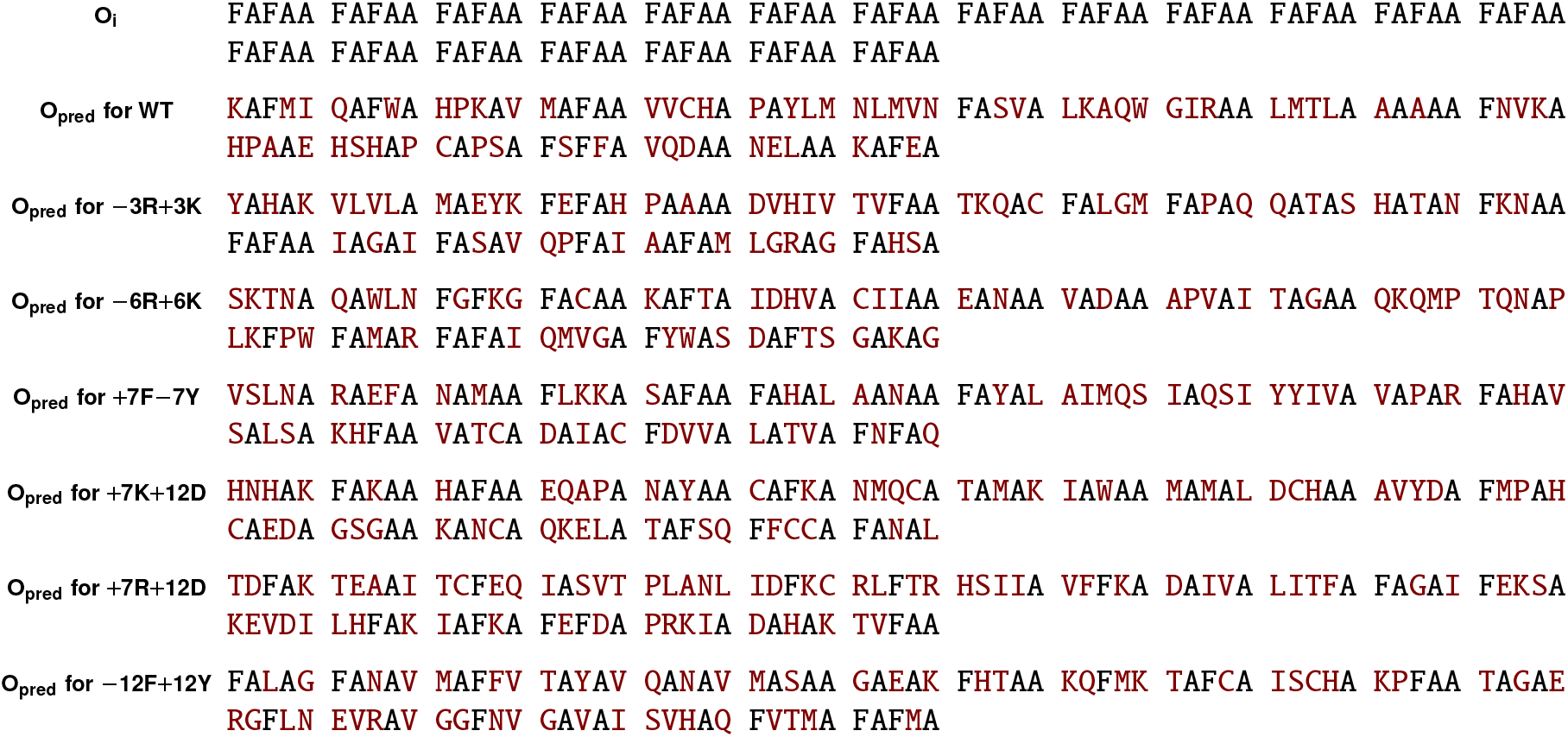

*Evolution of the inner protein when A1 LCD is on the outside.* In this case, I = (FAFAA)_20_ and O = (FIQII)_27_.

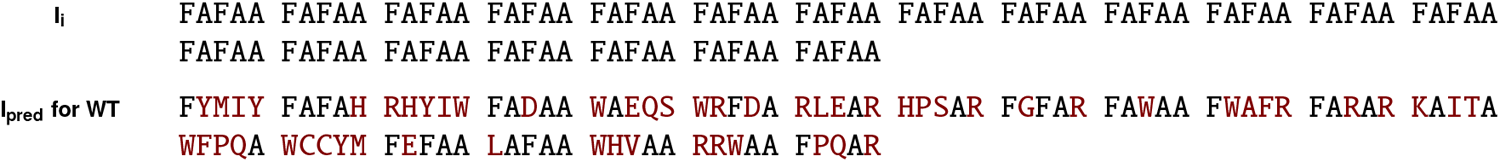

#### S10.13.3 Figure 3

*Evolution with partner protein fixed (Fig. 3a,b)*. Here, the protein O_i_ is (FAFAA)_5_. The protein I_i_ is F_50_ with added random mutations with a probability of 0.6.

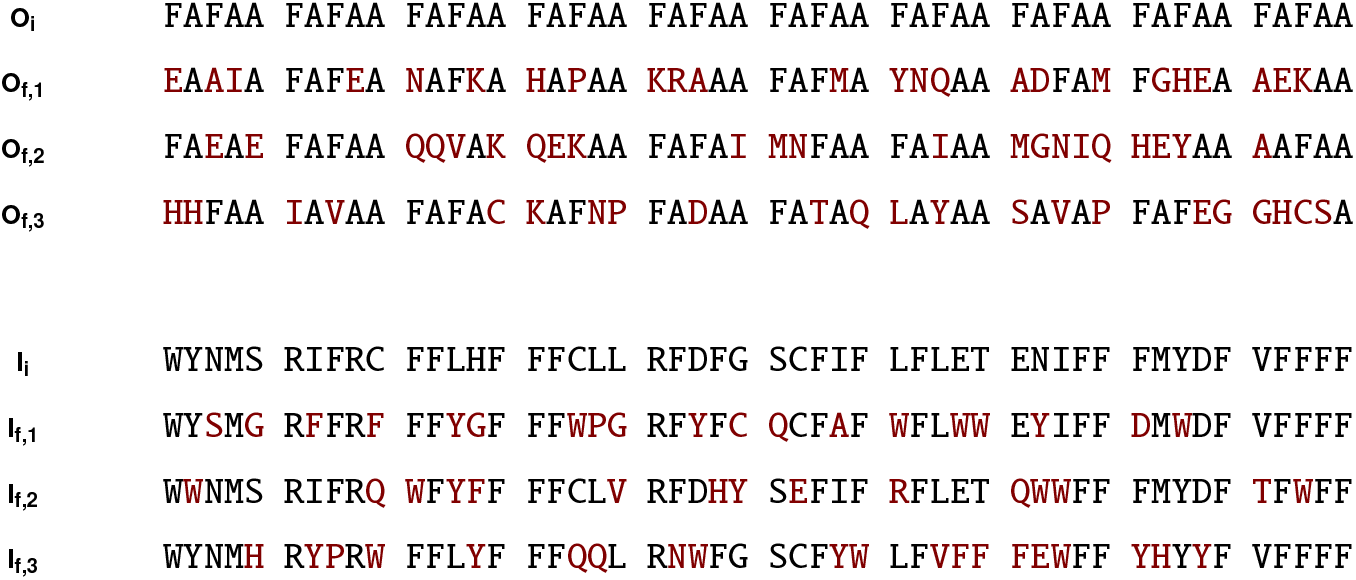

*Coevolution with A1 LCD (Fig. 3c,d).* In coevolution runs where A1 LCD ends up at the centre, we start with I = F_135_ and O = (FAFAA)_20_.

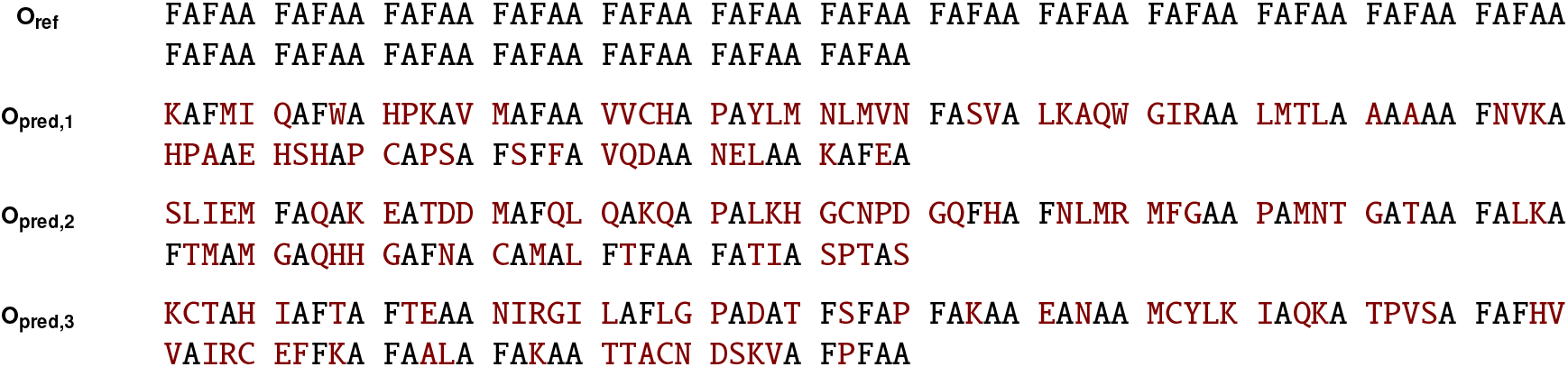

In coevolution runs where A1 LCD ends up on the outside, we start with I = (FAFAA)_20_ and O = (FIQII)_27_.

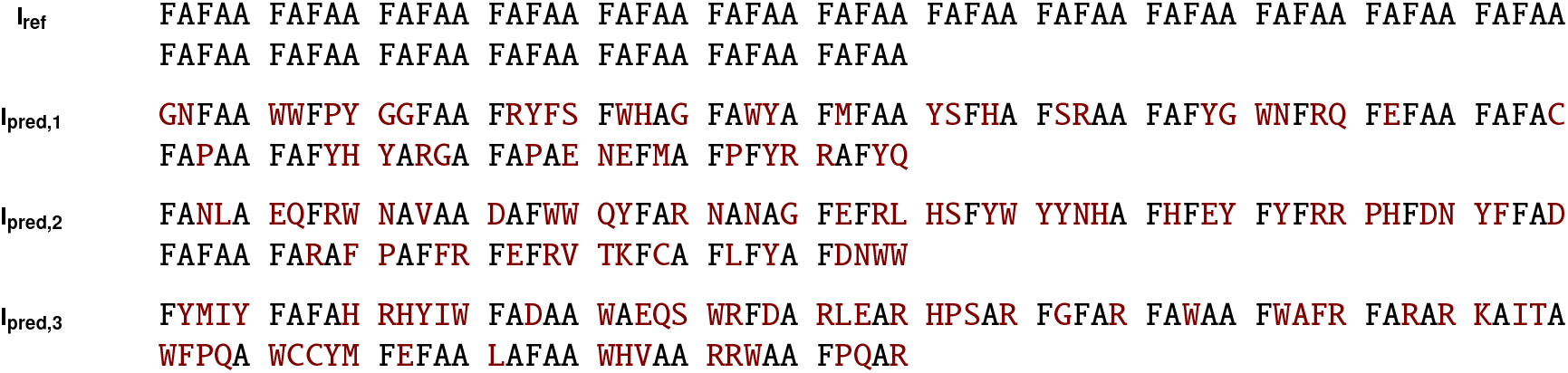

#### S10.13.4 Figure 5

For the case where spacers are shuffled, the amino-acid residues different from the reference evolved protein are highlighted in green.

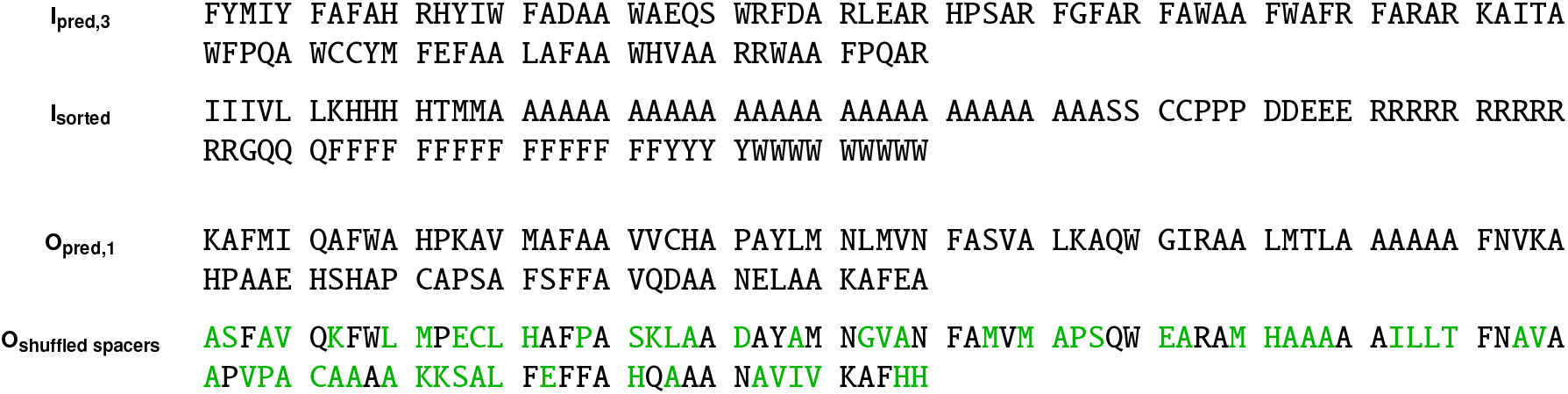

## Notes

### Competing Interest Statement

The authors have declared no competing interest.

